# From Networks to Traveling Waves and Back: Re-emergent Patterns Across Collective Phases in *C. elegans*

**DOI:** 10.64898/2026.06.02.729596

**Authors:** Assaf Pertzelan, Surabhi Sudevan, Gopika Ranjith, Siyu Serena Ding

**Affiliations:** Research Group Genes and Behavior, Max Planck Institute of Animal Behavior, 78464 Konstanz, Germany; International Max Planck Research School for Quantitative Behavior, Ecology and Evolution, 78464 Konstanz, Germany; Centre for the Advanced Study of Collective Behaviour, 78464 Konstanz, Germany

## Abstract

Thousands of *Caenorhabditis elegans* placed adjacent to a food patch self-organize sequentially into a visible branching network off food and then a coherent traveling consumption wave, a multi-phase collective phenomenon we term *wurmuration*. Dynamical networks and traveling waves have previously only been described separately in this organism. Using an agent-based model, we show that a single generative program produces both collective phases, and that what changes between phases is not the underlying rules but the food environment they act on. The program has four components: body elongation, roaming-dwelling switching, preferred local worm density navigation, and food chemotaxis. Each component is necessary within the model, and their combination reproduces the phenomenon across a range of parameters. We show that a cryptic network (revealed only by temporal integration) resembling the branching arrangement of the network phase re-emerges in the qualitatively distinct wave phase. Finally, tuning preferred density alone recapitulates N2 vs. *npr-1* differences in wave dynamics; N2 experiments additionally show greater variability between replicates in the cryptic network than *npr-1*. Our findings suggest that complex multi-phase collective phenomena can be unified by a shared generative program, shifting the description of spatial organization from the emergent patterns to their underlying components.

## Introduction

The same local interaction rules can produce qualitatively distinct macroscale collective patterns, a principle documented across biological scales and taxa^1–3^. In *Dictyostelium*, the same cAMP signaling produces sporadic, independent pulsing of cells at low density and synchronized oscillations that organize aggregation above a threshold density^4^; in desert locusts, local alignment rules produce disordered movement at low density and coherent directed marching at high density^5^; in fish schools, the same interaction rules produce polarized schooling, milling, and disordered swarming under shifting parameter regimes^6,7^. In *Caenorhabditis elegans*, collective patterns emerge from the interplay between population density, food availability, and ambient oxygen levels^8–12^, producing a repertoire of collective behaviors including stationary dense-cluster aggregation^11,13^, motile collective swarming^8,9,14,15^, dynamical network remodeling^12^, and density- and food-dependent pattern transitions^8^. Ding, Schumacher et al.^9^ showed that density-dependent speed switching, reversals and neighbor-taxis can reproduce both static aggregation and motile swarming from this minimal rule set, establishing that a shared generative program underlies these collective behaviors. While collective swarming has been reported across *C. elegans* population scales^8,9,14,15^, it has been modeled at the level of individuals only in small populations^9^. At large population scales, *C. elegans* collective behavior exhibits a fundamentally distinct phenomenology, encompassing multiple collective patterns within a single behavioral episode. No individual-level account exists to describe the emergence of and the transition between these patterns, nor to unify their relationship with previously described aggregation, swarming, and network remodeling.

Here we formally describe *wurmuration*, a large-scale, multi-phase collective phenomenon in *C. elegans*. When thousands of adult worms are deposited adjacent to a bacterial food patch, they first self-organize into a visible branching network of worm bundles in the off-food reservoir (network phase), then transition to a coherent traveling consumption wave (wave phase), in which no network is obvious from instantaneous observation. Temporal integration of worm positions, however, reveals that a cryptic network resembling the branching arrangement of the network phase re-emerges during the wave phase (Box 1). Both phases of wurmuration are purely emergent; no physical template connects them. Dynamical networks^12^ and collective swarming^8,15^ have each been documented in *C. elegans*, but their sequential emergence as phases of the same behavioral episode, and the generative basis of their relationship had not been established.

At the scale of thousands of animals, collective patterns are often described with continuum models, in which coarse-grained density and flux equations reproduce aggregation and traveling bands without resolving individuals. Keller and Segel^16^ treated the traveling bands of chemotactic bacteria described by Adler^17^ as driven by chemotaxis and consumption, and hydrodynamic theories extended the approach to flocking and to motility-induced phase separation^18,19^. In *C. elegans*, oxygen-dependent slowing within dense aggregates on a bacterial lawn has been described in these terms as motility-induced phase separation^8^. Such models describe patterns through population densities; however, to ask whether both wurmuration phases arise from the same rules followed by individual worms, the individuals themselves need to be modeled.

By developing an agent-based model, we show that four components suffice to produce both wurmuration phases. These components are body elongation, roaming-dwelling switching, navigation toward a preferred local worm density (*PD*, the density at which agents dwell most stably), and food chemotaxis; we show that each component is required within the model. A single between-strain parameter, *PD*, recapitulates the direction of the observed differences in wave dynamics between the laboratory reference strain N2 and the *npr-1* mutant DA609 [*npr-1*(*ad609*)X], hereafter referred to as *npr-1*. Our wurmuration model reproduces the cryptic network during the wave phase simply as a mechanical consequence of the same four components acting on the changed environment, with no structure carried over from the network phase. Together, these results suggest that the relevant level of description for the multi-phase wurmuration phenomenon lies in the shared generative components rather than in the emergent patterns they produce.

### Box 1 | Glossary of terms used in this study

#### Phases and patterns

**Wurmuration**: a **network phase** followed by a **wave phase**. A **phase** is a macroscopically distinct collective behavior, in the biological sense.

**Network**: a branching spatial arrangement of worm bundles. A **visible network** is seen in single frames during the network phase, off food. A **cryptic network** is revealed by the TCI during the wave phase, on food.

**Front**: the band of feeding worms at a given moment, bounded by a **leading edge** and a trailing edge.

**Wave**: a single front over time, advancing at near-constant speed and with a near-stationary profile.

#### Quantifications

**Wave metrics**: quantified by **wave speed** (the distance between the wave’s start and end positions divided by the time between them) and **wave bulkiness** (the maximal inscribed radius of the front, averaged over the wave phase).

**Front speed**: the slope of the leading edge in kymographs of the raw videos, an independent measure of the wave’s speed.

**Maximal inscribed radius**: the radius of the largest circle that fits inside a region.

**TCI** (temporal composite imprint): an image in which each pixel keeps the darkest value it reached over the recording, marking regions of peak worm occupancy.

**Coverage geometry**: the arrangement of **bundles** (dark regions that worms visited) and **holes** (bright enclosed regions they left unvisited) in a TCI, quantified by **large hole area** (the area of the fifth-largest hole) and **TCI bulkiness** (the maximal inscribed radius of the bundles).

#### Model

**Generative program**: the four model components shared by both phases, namely body elongation, roaming-dwelling switching, navigation toward a preferred local worm density (**PD**, the density at which agents dwell most stably), and food chemotaxis.

**Leave-one-out simulations**: each component removed in turn.

**No-network-phase control**: the wave starts from a **structureless front** made of agents seeded randomly in a narrow strip spanning the initial 5% of the food patch, with no off-food network phase.

**Sensitivity analysis**: six parameters changed by ±20%, one at a time; each change is a perturbation.

**Trade-off**: across perturbations in the model, larger holes come with thinner bundles.

## Results

### Wurmuration exhibits multi-phase spatiotemporal dynamics

To characterize wurmuration, we deposited ∼20,000 synchronized Day 1 *C. elegans* adults in a concentrated droplet adjacent to a ∼110 x 30 mm rectangular *E. coli* OP50 lawn (Fig. 1a). Time-lapse imaging revealed a stereotyped progression of collective dynamics (Fig. 1b; Supplementary Movie 1a, c). Initially, worms formed a visible network within the off-food worm reservoir as the liquid absorbed into the hydrogel media (network phase) (Fig. 1b-ii). ’Network’ in our study refers to a branching spatial arrangement of worm bundles, not to a network of interactions between individuals. The visible network subsequently drained toward the food patch, giving rise to a coherent traveling wave advancing across the bacterial lawn (wave phase) (Fig. 1b-iii,iv). ’Wave’ here refers to a single consumption front over time; the ‘front’ is the band of feeding worms at any given moment, bounded by a leading edge and a trailing edge. Kymographs of the raw videos show that the front advances across the lawn at an approximately constant speed and with a near-stationary profile (Supplementary Fig. 1), meeting the two defining features of a traveling wave in biology^16,20^. Properties of periodic waves such as wavelength and period do not apply in our case. Instead, we quantify the wave by its speed (*wave speed*) and by the thickness of the front, measured as its maximal inscribed radius, the radius of the largest circle that fits inside the front (*wave bulkiness*). We use the term ’phase’ throughout in the biological sense to refer to macroscopically distinct collective behaviors, rather than in the thermodynamic sense of an order-parameter transition. Close observation of wave phase dynamics revealed that the front did not advance uniformly across the food patch: worms appeared to preferentially follow along certain routes, bundling around discrete areas rather than passing through them, leaving those areas progressively encircled but unvisited as the front advanced, a hint of spatial patterning during the wave phase (Supplementary Movie 2).

**Figure 1.**
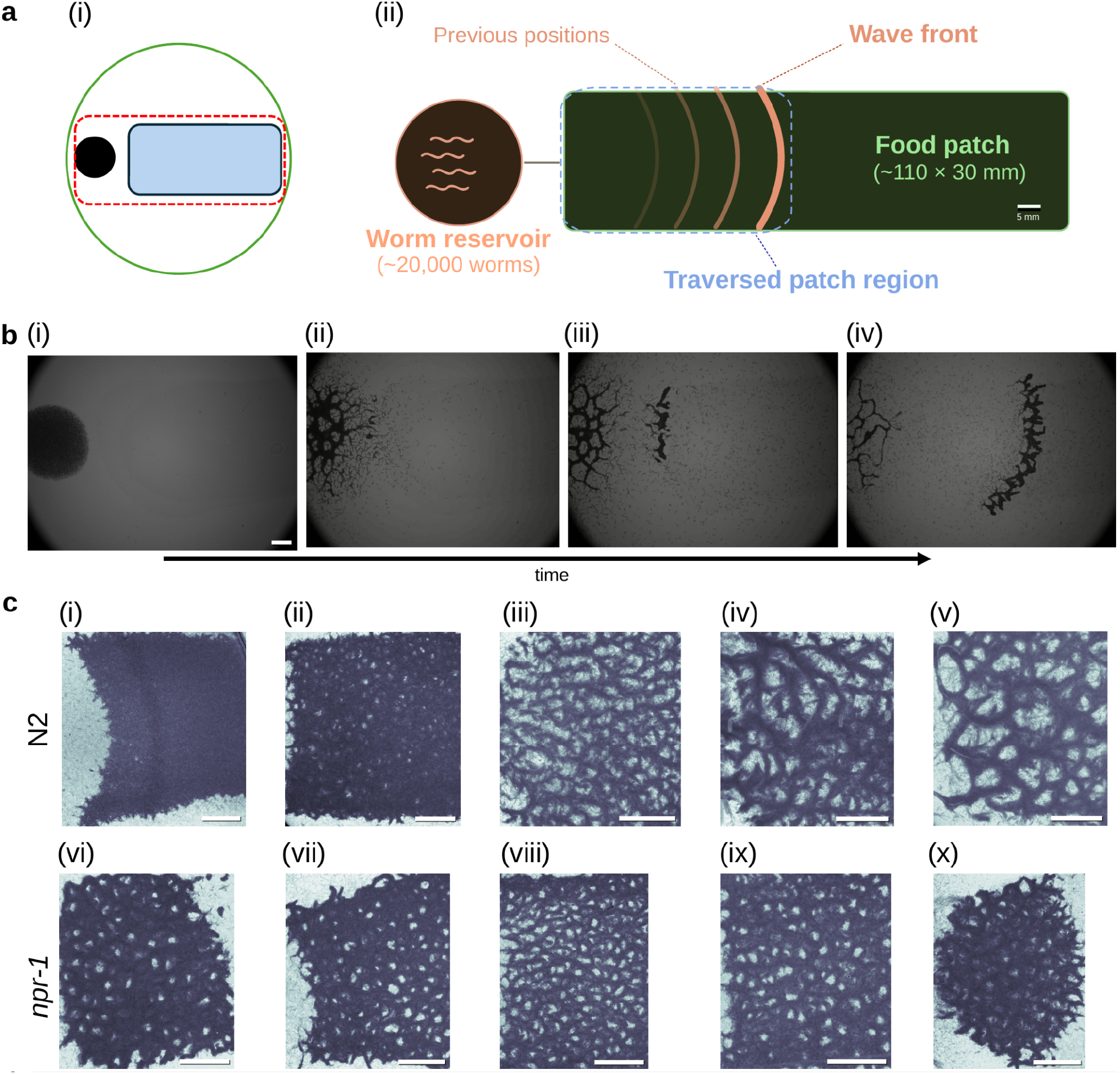
Wurmuration exhibits a visible network and a cryptic network across its two phases. **a** (i) Schematic of the experimental setup in the 150 mm-diameter behavioral arena. The worm reservoir (black circle) is positioned at the left edge of the bacterial lawn (blue rounded rectangle) inside the Petri dish (green circle), with the detailed view area indicated by the dashed red rounded rectangle. Not drawn to scale. (ii) Detailed view of the interaction area. Individuals may navigate inside the Petri dish beyond the boundaries of this view. **b** Time-lapse sequence (*npr-1*) showing: (i) initial reservoir, (ii) network phase: visible branching network in the worm reservoir, (iii) network-to-wave transition: visible network draining and wave emergence, (iv) wave phase: traveling wave advancing across the bacterial lawn. **c** Representative TCIs showing the cryptic network during the wave phase. Dark bundles mark regions that worms visited; bright holes are enclosed regions they left unvisited. (i–v) N2 strain showing variable coverage geometry across replicates, from thick bundles with few holes to thin bundles around large holes. (vi–x) *npr-1* showing more consistent coverage geometry and hole size distribution across replicates. All scale bars: 5 mm.

To reveal worm positions across the entire wave phase we generated images in which each pixel retains the darkest value recorded at that location across all frames, marking regions of peak worm occupancy. These temporal composite imprints (TCIs; Fig. 1c) displayed a distinctive cryptic network, with dark bundles where worms visited and bright enclosed holes that they left unvisited. We use the term ‘coverage geometry’ to refer to the spatial arrangement of bundles and holes in a TCI, and quantify it by the area of the fifth-largest hole (*large hole area*) and the thickness of the bundles, measured as the maximal inscribed radius within them (*TCI bulkiness*; Methods). This cryptic network, invisible in instantaneous frames but recoverable through temporal integration, was observed robustly in both N2 and *npr-1*. N2 TCIs (Fig. 1c, i–v) showed greater variability between experimental replicates in coverage geometry compared to *npr-1* (Fig. 1c, vi–x). The cryptic network was further observed across a broad range of strain genetics, population compositions, arena sizes, and food conditions in independent experiments not pooled with the main dataset (Supplementary Fig. 2), indicating that networks are a general feature of wurmuration, both as a visible network in the off-food network phase and as a cryptic network in the on-food wave phase.

### Agent-based simulations recapitulate experimental multi-phase dynamics

We developed an agent-based model to investigate the minimal requirements for the observed collective dynamics. In the model, self-propelled agents representing individual worms (Fig. 2a,b; Supplementary Movie 3) are initialized in a dense off-food reservoir adjacent to a rectangular food patch. Each agent senses three quantities: the local worm density along its own body and around its head, the magnitude of a food odor field at its head, and the local food concentration at its head. Agents do not interact directly, have no access to neighbor identity, alignment or attraction, and affect one another only by adding to the local density and by depleting food (Supplementary Fig. 3). Agents switch between roaming and dwelling with probabilities set by the density mismatch summed along the whole body, and a roaming agent that is not odor-guided steps toward whichever neighboring site brings it closest to *PD*, with crowding above *PD* penalized three times as heavily as the same shortfall below (Supplementary Methods, Agent-Based Simulation). The probability of redirecting toward the food instead rises with odor magnitude and falls with local food concentration. Chemotaxis is therefore suppressed on intact food, where agents accumulate until local worm density reaches *PD* and then dwell while feeding. As this food is depleted, chemotaxis resumes and draws them toward the intact food ahead, so the front advances (Supplementary Methods, Numerical illustration of component interactions).

**Figure 2.**
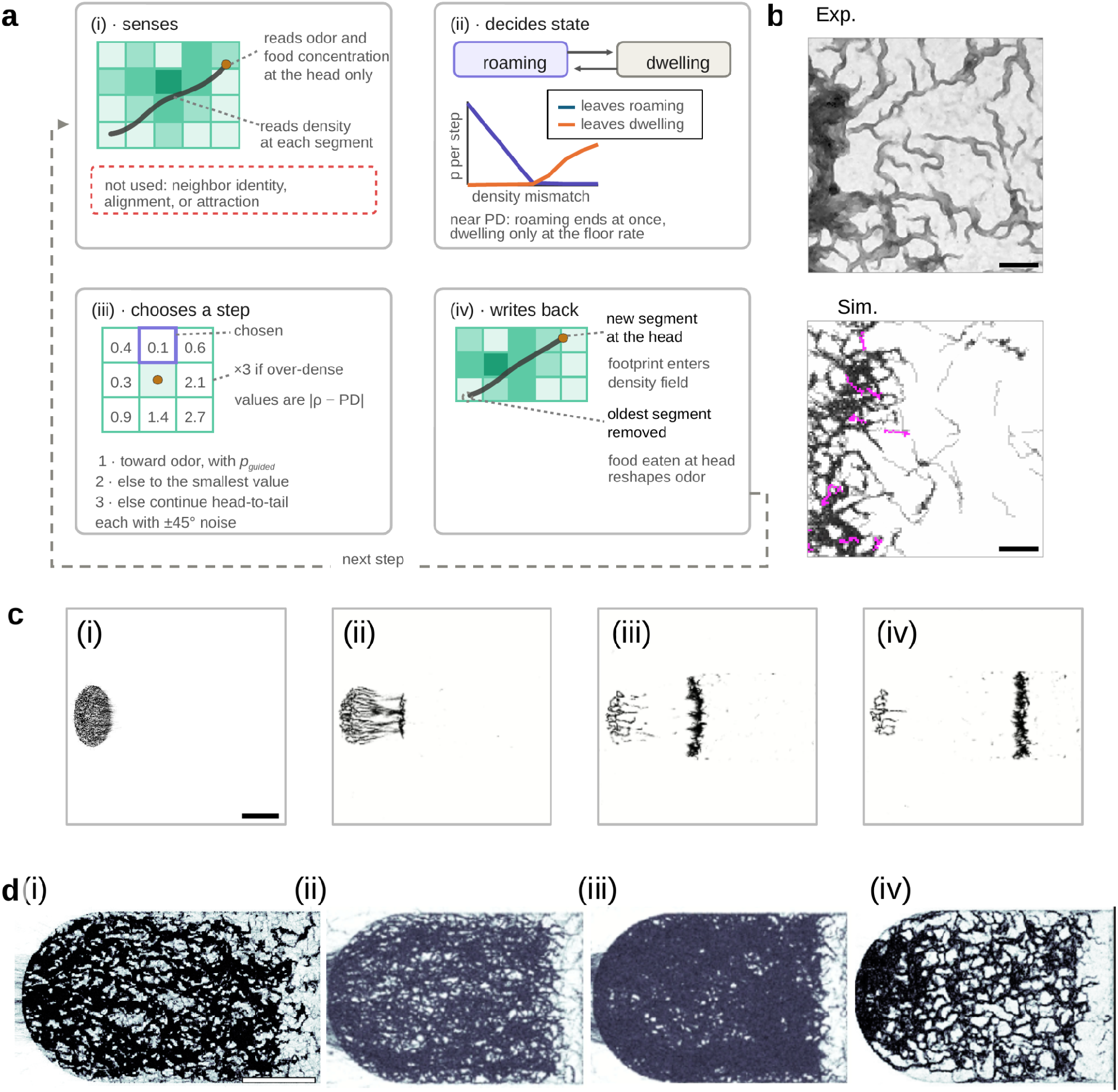
Agent-based simulations recapitulate experimental multi-phase dynamics. **a** Schematic of one agent in one time step (steps i–iv); in (ii), the probabilities per step of leaving roaming (blue line) and dwelling (orange line) are plotted against the density mismatch summed along the body (zero at *PD*) ; ρ, local worm density; *p*_guided_, probability of an odor-guided step; see Supplementary Methods, Agent-Based Simulation, for the full description. **b** Close-up views of the front at matched physical scale for experiment (N2) and simulation (*PD* = 6; rendering in Methods, Simulation Analysis). A random 1% of simulated agents are shown in magenta. The experimental frame is shown with a gamma of 0.6, edge-preserving smoothing and unsharp masking. Scale bars: 1 mm. Also see Supplementary Movie 3. **c** Replication of the observed wurmuration dynamics from visible network formation in the worm reservoir to traveling wave initiation and advance. Scale bar: 5 mm. Compare to Fig. 1b. **d** Multiple examples of TCIs of simulated agent positions over full simulation, showing consistent occurrence of the cryptic network across different parameterizations. Scale bar: 5 mm. Compare to experimental TCIs in Fig. 1c.

The model successfully recapitulated the complete wurmuration sequence from the network phase through the emergence and advance of the traveling wave (Fig. 2c, compare to Fig. 1b; side-by-side comparison in Supplementary Movie 1). The simulations also reproduced the cryptic network observed in experimental TCIs (Fig. 2d, compare to Fig. 1c); quantitative comparison between experiments and simulations will follow in subsequent sections. A cryptic network also emerges in a no-network-phase control, in which the wave starts from a structureless front made of agents seeded randomly in a narrow strip spanning the initial 5% of the food patch (Supplementary Fig. 4), so the wave phase can independently generate cryptic networks without a preceding off-food reservoir or visible network.

To test model robustness in a sensitivity analysis, we varied body length, the baseline roaming and dwelling bouts, odor threshold, angular noise and collective feeding efficiency, one parameter at a time, by ±20% with 40 replicates per condition. The wave and its cryptic network formed under every parameter perturbation, and no perturbation shifted the mean of any wave or coverage geometry metric by more than 39% from its default value (Supplementary Fig. 5; Supplementary Table 1). The wave and its cryptic network also formed at the two *PD* values used for the simulated strains, which differ twofold (next section). The four components are therefore sufficient to produce the phenomenology across a region of parameter space, and not only at a single tuned point.

We also conducted leave-one-out simulations to test whether each of the four components is necessary within this model architecture and parameterization (Fig. 3; Supplementary Movie 4; Supplementary Methods, Leave-one-out simulations). Body elongation generates nematic order, as two agents alone sit far below *PD* so each gains by overlapping the other (agents can occupy the same sites), and elongated bodies overlap most when their long axes are parallel. Since both agents keep moving, they then travel together or in opposite directions, generating nematic rather than polar order. Elongated agents became strongly aligned after pairwise interactions while round agents did not (mean resultant length *r* = 0.11 before and 0.45 after interaction for elongated agents, *n* = 868 interactions; *r* = 0.20 before and 0.15 after for round agents, *n* = 1,267; Supplementary Table 2), and without elongation the patch was consumed in a uniform advancing front with no cryptic network (Fig. 3a,b). Roaming-dwelling switching is essential for sustaining the cryptic network: the agents generate the structure while roaming and the switch to dwelling keeps it stable; roaming-only agents produce transient fronts that subsequently fragmented (Fig. 3c). *PD* navigation, where agents settle near moderate local crowding and disperse from both local density extremes, is required for bundle formation; removing it leaves no cryptic network (Fig. 3d). Food chemotaxis converts density-driven dynamics into a traveling wave; without it, agents form continuously remodeling visible networks but no coherent front (Fig. 3e), resembling the dynamic network reported by Sugi et al.^12^. Experimental worm populations deposited without food also formed visible networks but no front (Supplementary Fig. 6). Together, these four model components produce the cryptic network through their interaction with local food depletion.

**Figure 3.**
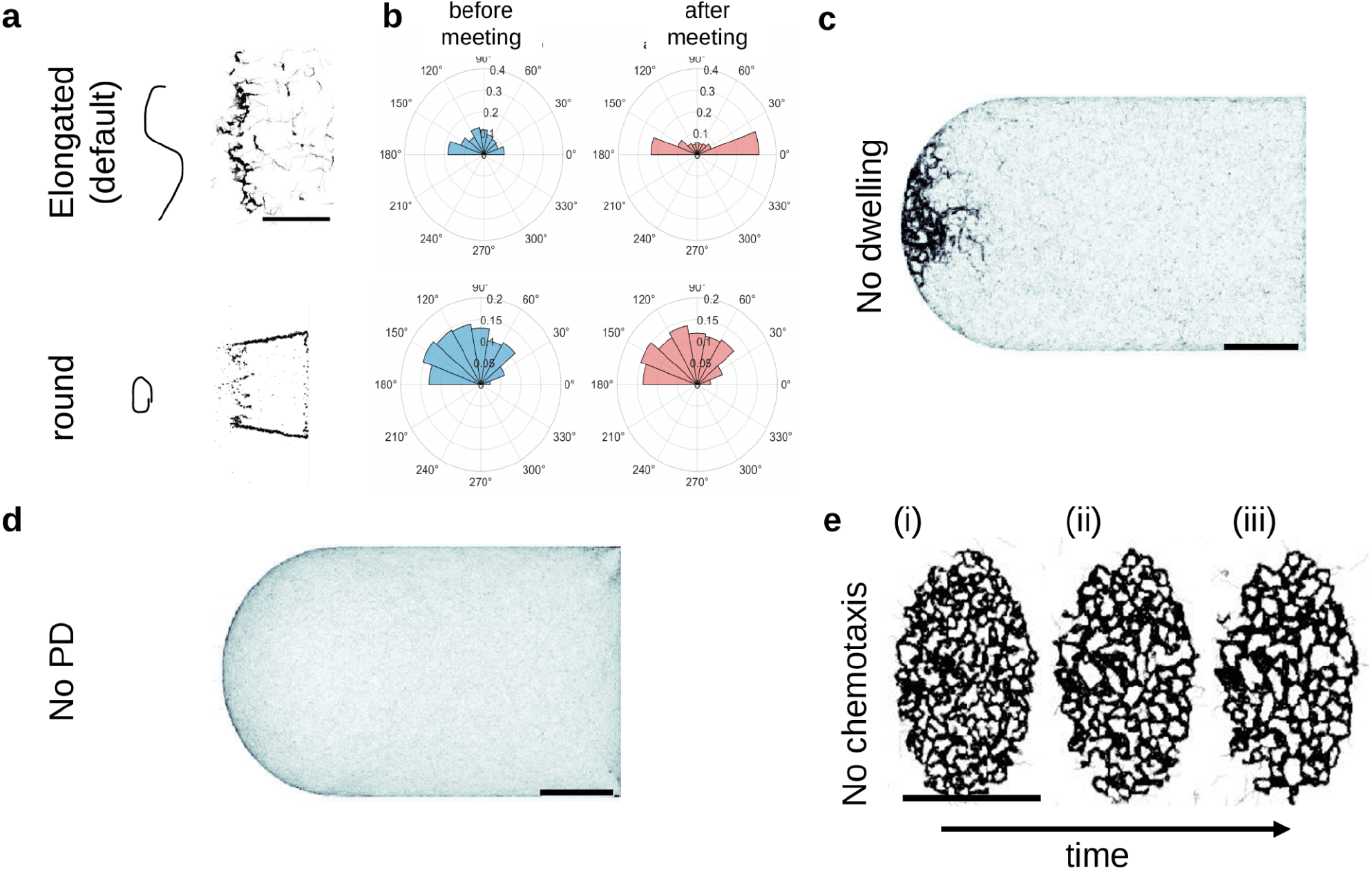
Necessity of each model component. See Supplementary Movie 4 and Supplementary Methods, Leave-one-out simulations. **a** Snapshots of elongated vs. round agents. **b** Angular distribution of pairwise approach and departure angles, corrected for geometric sampling bias (see Methods, Simulation Analysis; Supplementary Methods, Agent-Based Simulation, Nematic alignment analysis). No explicit alignment rule is implemented; nematic order emerges from body elongation. **c** TCI, roaming-only agents (no dwelling). **d** TCI, no *PD* navigation. **e** Time-lapse, no food chemotaxis. All scale bars: 5 mm. Source data are provided as a Source Data file.

### Varying *PD* captures strain differences in wave metrics but not in coverage geometry

Having established that the general wurmuration behavioral sequence emerges from a minimal set of components, we next asked what parameter variation accounts for the observed differences between strains. *npr-1* animals preferentially aggregate at higher local worm densities, partly mediated by a preference for the local low-oxygen microenvironments created by conspecific respiration and limited oxygen diffusion in dense groups^10,11,13,21^. In our model, this enhanced density-seeking tendency is represented by a higher *PD*, the local worm density at which agents dwell stably. We therefore set *PD* = 6 for N2 and *PD* = 12 for *npr-1* with all other parameters held constant, reflecting the relative difference in density-seeking tendency between strains, to test whether this single parameter variation explains the strain differences (Supplementary Movie 1, panels a–d). The simulations are intended to reproduce the patterns and the direction of strain differences rather than absolute values, which differ substantially in magnitude from the experiments (Supplementary Note 1).

This single *PD* difference recapitulated the observed strain differences in wave metrics (Fig. 4a; Supplementary Table 2). In both experimental data and simulation sets, N2 *wave speed* is higher than *npr-1* (Hedges’ *g* = 1.30, *P* = 0.012 in experiments; *g* = 0.82, *P* = 0.011 in simulations), and N2 fronts are broader than *npr-1* as measured by *wave bulkiness* (Fig. 4b-i, yellow circles; *g* = 1.08, *P* = 0.041 in experiments; *g* = 4.01, *P* < 0.001 in simulations). Front speed, measured independently as the slope of the leading edge in the kymographs (Fig. 4b-ii; Supplementary Fig. 1), closely matches *wave speed* in the experiments (Spearman’s ρ = 0.93 for N2, ρ = 0.90 for *npr-1*). The agreement in *wave speed* and *wave bulkiness* across both experimental data and simulation sets therefore supports the interpretation that varying *PD* alone captures the direction of the strain differences in wave metrics.

**Figure 4.**
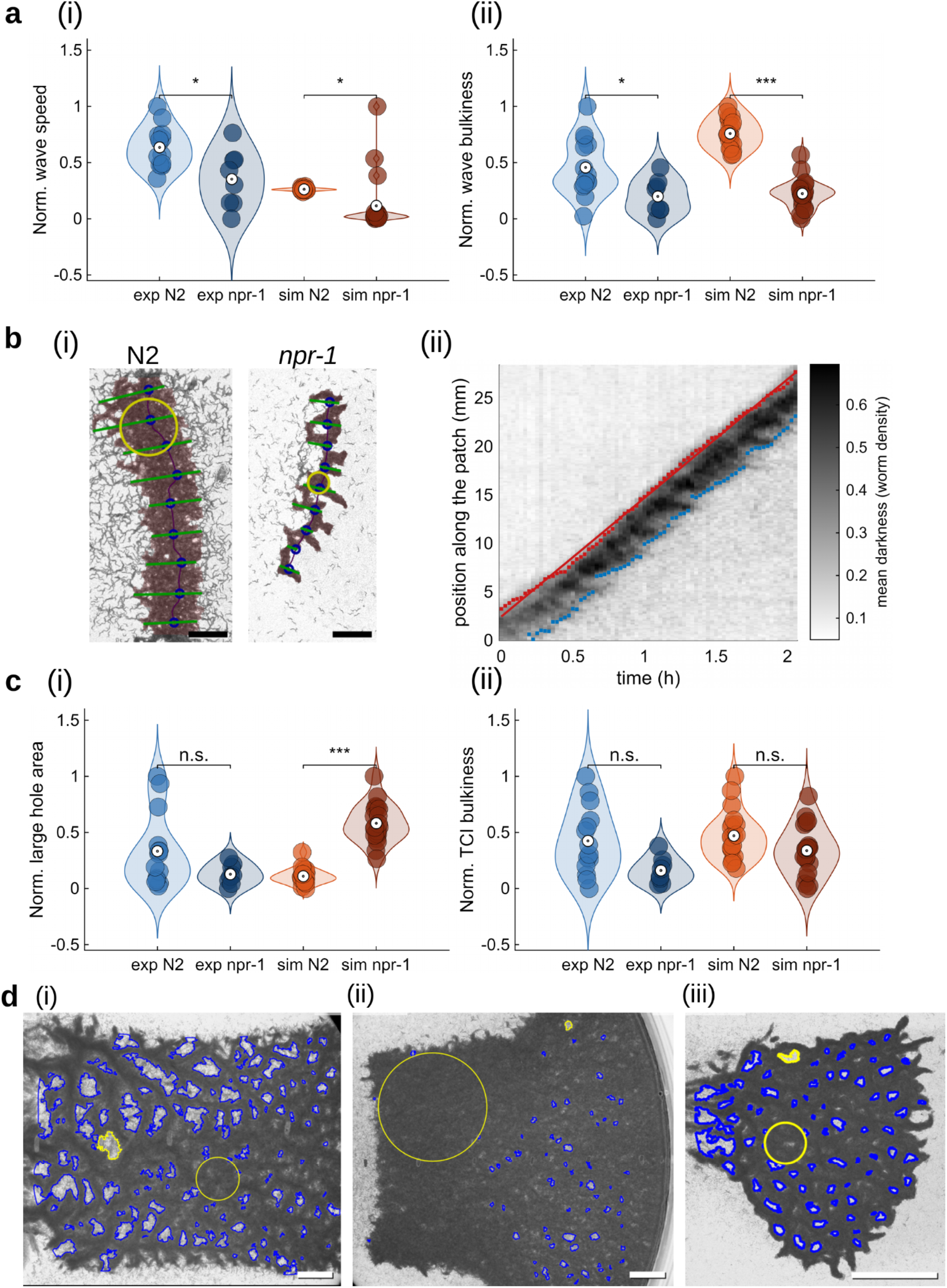
Varying *PD* recapitulates strain differences in wave metrics but not coverage geometry. **a** Wave metrics in experiments (blue) and simulations (orange). Simulated N2 (*PD* = 6) and *npr-1* (*PD* = 12) differ only in *PD*. Values are independently min–max normalized to [0–1] over the combined N2 and *npr-1* range within experiments and within simulations for visual comparison only; the normalization leaves *P* values unchanged; (i) *wave speed*, (ii) *wave bulkiness*. Points are replicates; white circles mark the means. Tests are two-sided permutation tests; stars show Bonferroni-adjusted *P* values. *: *P* < 0.05; ***: *P* < 0.001. Experimental sample sizes are: N2 *n* = 14, *npr-1 n* = 8; simulation sample sizes are: N2 *n* = 20, *npr-1 n* = 20. **b** (i) Representative fronts of an N2 and an *npr-1* wave. Yellow circles indicate *wave bulkiness*; green lines show sampled cross-sections of the front (see Supplementary Methods, Wave Metric Extraction). Scale bars: 5 mm. (ii) Sample kymograph of the *npr-1* experiment in Supplementary Fig. 1a-ii, with the detected leading edge (red points), its linear fit (red line; 12.24 mm h^-1^, *R*² = 0.997) and the trailing edge (blue points) of the front. **c** Coverage geometry metrics in experiments (blue) and simulations (orange). Values are independently min–max normalized to [0–1] over the combined N2 and *npr-1* range within experiments and within simulations for visual comparison only; the normalization leaves *P* values unchanged; (i) *large hole area*, (ii) *TCI bulkiness*. ***: *P* < 0.001; n.s.: not significant. Experimental sample sizes are: N2 *n* = 14, *npr-1 n* = 8; simulation sample sizes are: N2 *n* = 20, *npr-1 n* = 20. **d** Representative experimental TCIs: (i) N2 with high *large hole area* and low *TCI bulkiness*, (ii) N2 with low *large hole area* and high *TCI bulkiness*, (iii) *npr-1*. Yellow regions are the 5th largest hole and the bulkiness radius. Scale bars: 5 mm. Source data are provided as a Source Data file.

However, varying *PD* alone was insufficient to reproduce the observed differences in coverage geometry between strains (Fig. 4c; Supplementary Table 2; Supplementary Methods, Coverage Geometry Extraction). For *large hole area,* the experiments show no significant strain difference (*g* = 0.73, *P* = 0.201) whereas the simulations show higher values for *npr-1* than N2 (*g* = −3.52, *P* < 0.001). Experimentally the strains differ in spread more than in mean, with pronounced variability between replicates in N2 but not in *npr-1*, which the model does not capture. Neither the experiments nor the simulations show a significant strain difference in *TCI bulkiness* (*g* = 0.99, *P* = 0.058 in experiments; *g* = 0.59, *P* = 0.123 in simulations). Figure 4d shows representative experimental TCIs illustrating different combinations of the two coverage geometry metrics: N2 with high *large hole area* and low *TCI bulkiness* (Fig. 4d-i), N2 with low *large hole area* and high *TCI bulkiness* (Fig. 4d-ii), and *npr-1* (Fig. 4d-iii). We further characterized the variability between N2 experimental replicates in the following section.

### Variability between replicates in coverage geometry is consistent with a trade-off in the model

N2 experiments show substantially greater variability between replicates in coverage geometry than *npr-1* experiments. Standard deviations are 3.5-fold higher in N2 than *npr-1* for *large hole area* (*P* = 0.033) and 2.6-fold higher for *TCI bulkiness* (*P* = 0.021; permutation tests on the standard deviation ratio; Supplementary Table 2; see Methods); *npr-1* experiments correspondingly occupy a tighter region of this two-metric space compared to N2 (Fig. 5a-i). In both strains, experiments with higher *large hole area* show a trend toward thinner bundles (Spearman’s correlation; N2 ρ = −0.53, *P* = 0.057; *npr-1* ρ = −0.62, *P* = 0.115). By contrast, simulations run at fixed parameters failed to reproduce this pattern, as their scatter is uncorrelated (ρ = 0.20, *P* = 0.404 in N2) and shows no greater variability in N2. The standard deviation ratios between N2 and *npr-1* simulations are 0.42 for *large hole area* and 0.98 for *TCI bulkiness* (Fig. 5a-ii), so variability between simulation replicates alone does not generate the observed experimental N2 spread.

**Figure 5.**
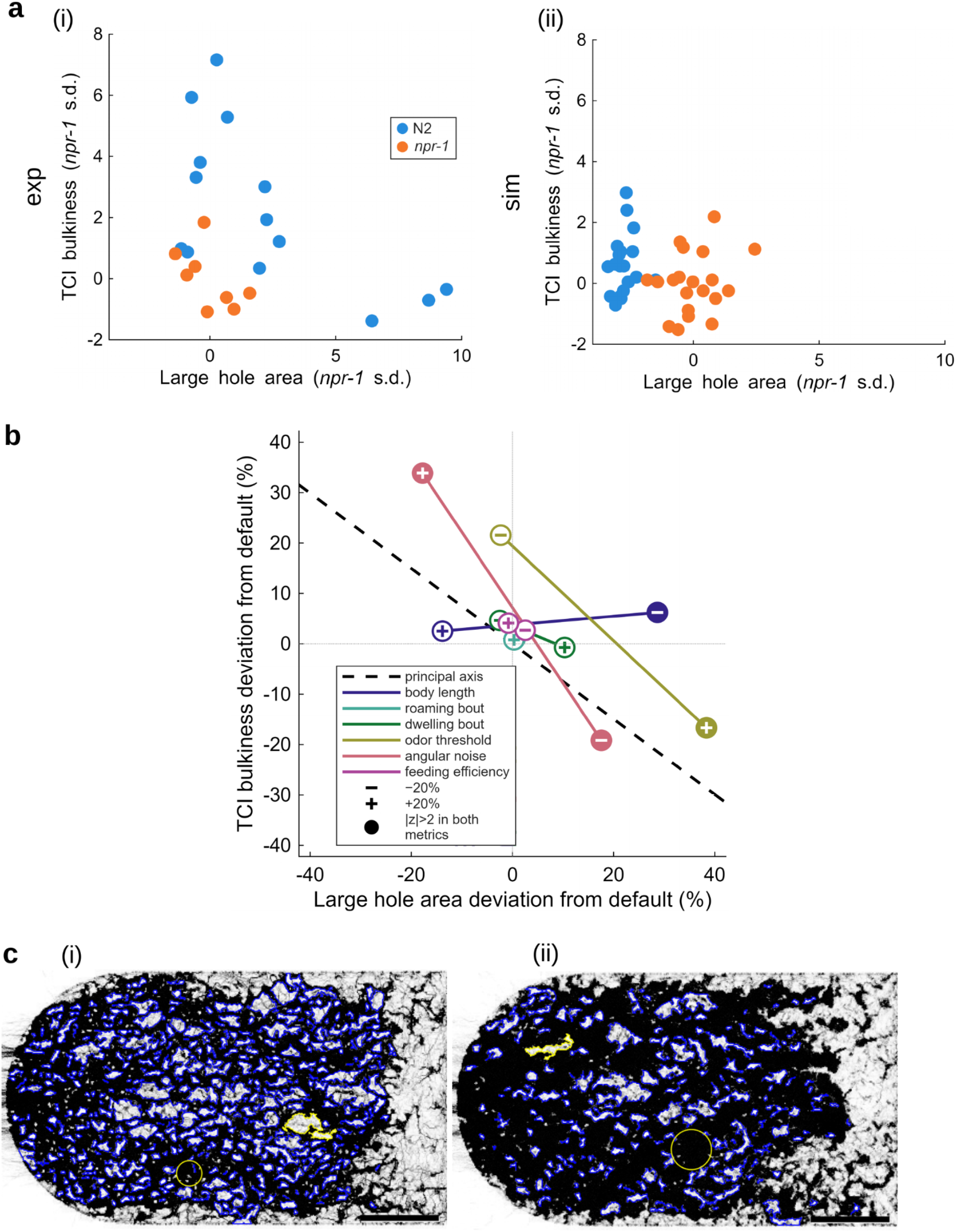
Pronounced variability between N2 experimental replicates in coverage geometry is consistent with a trade-off in the model. **a** Scatter of *large hole area* vs. *TCI bulkiness* for (i) experiments and (ii) simulations at fixed *PD*s. Each axis shows deviations from the *npr-1* mean in units of the *npr-1* s.d. of the same panel. N2 standard deviation is ∼3.5-fold that of *npr-1* for *large hole area* (*P* = 0.033) and ∼2.6-fold for *TCI bulkiness* (*P* = 0.021); permutation *P* values are in Supplementary Table 2. **b** Response of the two coverage geometry metrics to ±20% perturbations of each parameter, applied one at a time, in the sensitivity analysis (Supplementary Fig. 5; *n* = 40 replicates per condition). Lines join each parameter’s −20% and +20% points (circles marked − and +, respectively); the dashed line is the principal axis through the origin (slope −0.75). Of the four perturbations with |*z*| > 2 in both metrics (filled markers; *z*, the shift divided by its standard error), all but body length −20% move them in opposite directions. **c** Representative simulation TCIs at the two ends of the principal axis, marked as in Fig. 4d: (i) angular noise −20%, (ii) angular noise +20%. Scale bars: 5 mm. The trends in a-i and b agree in direction rather than in magnitude (Supplementary Note 1). Source data are provided as a Source Data file.

In the sensitivity analysis, the two coverage geometry metrics move along a single axis rather than independently. Eight of the 12 perturbations move *large hole area* and *TCI bulkiness* in opposite directions, and a principal axis through the origin with a slope of −0.75 captures 78% of the total variation (Fig. 5b; Supplementary Fig. 5). Representative simulation TCIs at both ends of this axis are shown in Fig. 5c. This trade-off of larger holes with thinner bundles, is therefore a property of the model as a whole and not the signature of a single parameter.

The variability between N2 experiments in coverage geometry is consistent with this trade-off. Our results suggest that N2 populations may produce more variable coverage geometry along this trade-off, potentially due to a less consistent fraction of worms joining the collective movement, an explanation that we have not tested (Supplementary Note 2). *npr-1*, by contrast, produces more stereotyped coverage geometry across replicates, consistent with stronger sociality constraining collective outcomes.

## Discussion

We describe wurmuration, a large-scale multi-phase collective phenomenon in *C. elegans* in which the same population sequentially self-organizes into a visible branching network and then a coherent traveling consumption wave within a single behavioral episode. The sequential emergence of the network and wave phases, each previously described only in isolation in this organism^8,12,15^, constitutes our primary phenomenological finding. Temporal integration of the wave phase recovers a cryptic network hidden from instantaneous observations alone. These findings together call for a mechanistic account that can encompass both collective phases and their spatial relationship within a single framework. Within our model architecture and its parameterization, four components (body elongation, roaming-dwelling switching, *PD* navigation, and food chemotaxis) interact together to generate both wurmuration phases. What changes between the phases is not the components themselves but the environment they act on: food-absent in the network phase, food-present and depleting in the wave phase. The model reproduces the complete behavioral sequence, and the wave and its cryptic network form robustly in the sensitivity analysis (Supplementary Fig. 5). Reaching a better quantitative agreement between experiments and simulations (Supplementary Note 1) and experimentally testing model predictions (Supplementary Note 2) remain important future directions.

The strain comparison positions *PD* as a tunable parameter of wave dynamics. Varying *PD* as the single between-strain parameter recapitulates directional differences across both wave metrics, with higher *PD* producing slower, more compact waves in *npr-1* simulations than in N2; whether experimentally raising density preference in the worms produces the same effect remains to be tested (Supplementary Note 2). In coverage geometry, however, the simulations do not reproduce the experimental strain differences in their spread. N2 experiments show substantially greater variability between replicates in coverage geometry than *npr-1* that the simulations fail to capture, potentially suggesting a missing variable that shifts N2 replicates along the model’s trade-off. We hypothesize that the fraction of worms joining the collective movement could be a candidate variable and that the stronger sociality in *npr-1* constrains the movement along this trade-off to produce more consistent collective outcomes. However, this hypothesis remains untested and the actual biological source and mechanisms driving the variation between replicates in low-sociality N2 remain an open question for future investigation.

The density-centric logic underlying wurmuration connects with the framework established by Ding, Schumacher et al.^9^ across several orders of magnitude in collective scale. Where that work established that density-dependent speed state switching and neighbor-taxis reproduce aggregation and swarming in small populations, the same density-centric logic here encompasses large-population visible network formation and a traveling wave within a single behavioral episode. We further extended this previous work in two specific directions. First, the neighbor-taxis rule required for cohesion in that model is here subsumed into *PD* navigation alone, suggesting that movement toward preferred local worm density regions may be sufficient for both state-switching and directional cohesion without a separate attraction term. Second, while Ding, Schumacher et al. showed that local food depletion drives aggregate motility in the swarming phase, food chemotaxis in our model acts as an explicit active component converting density-driven visible network dynamics into a traveling consumption wave. Here, agents navigate toward food ahead of the wave rather than disbanding passively from depleted regions; the wave’s progressive depletion of local food thus sustains food gradient-directed collective advance without any change in the four components themselves. Our model does not specify what sets the *PD–*it could be a behavioral preference, humidity (as in Sugi et al.’s system^12^), or another cause. We did not test for the threshold worm density that Sugi et al. report for network formation.

Comparing the wurmuration model to other *C. elegans* collective behavior frameworks clarifies the specific mechanistic contribution of each component to network and wave formation. Chen and Ferrell^22^ showed that density-thresholded aggregation can arise from purely passive condensation: worms performing random walks slow upon colony contact, without any active taxis toward conspecifics producing density-dependent clustering. The spatial outcome is round, liquid-like aggregates, not branching networks, establishing that network formation requires active spatial patterning that passive velocity reduction alone cannot provide. Sugi et al.^12^ identified a candidate: density-dependent aggregation combined with nematic alignment generates branching dynamic networks in the absence of food. Critically, nematic ordering was an empirically observed property in their system and encoded as an explicit alignment rule in the model. In our wurmuration model, nematic organization is not an imposed fixed input but an emergent mechanistic output: multi-segment elongated worms with directional persistence generate bundle-like alignment under *PD* constraints without any explicit alignment interaction, as demonstrated by comparison with isotropic round agents (Fig. 3a,b). Our model thus identifies body geometry as sufficient for nematic order, a property that Sugi et al.’s model presupposes and Chen and Ferrell’s does not require. Food chemotaxis, absent from both prior models, then completes our model by converting the continuous remodeling of the visible network into a traveling wave within the same behavioral episode in wurmuration.

Our consumption wave is a single front over time of the kind described for traveling bands of chemotactic bacteria^16,17^, which continuum models treat at the level of averaged density fields. Here we ask instead which individual-level rules produce such a front; the coarse-grained limit of our model remains to be derived. The model also bears on motility-induced phase separation (MIPS), invoked previously for oxygen-dependent aggregation on a bacterial lawn^8^. In our model agents either roam at a fixed speed or dwell, so their effective speed (roaming speed times the fraction of time spent roaming) follows from the switching rule alone (Supplementary Methods, Numerical illustration of component interactions, Effective speed and bout durations). Below *PD*, effective speed falls with density as in MIPS, but the model differs from classical MIPS in two respects. First, effective speed here is non-monotonic rather than monotonically decreasing with density. It is minimal at *PD* and rises steeply above it, so the density inside aggregates is capped near *PD* by the rule itself. Secondly, in our model agents choose their direction from the local density (*PD* navigation), whereas in classical MIPS the direction of motion is independent of density and aggregation follows from speed reduction alone^18^.

The model also shows how a cryptic network can re-emerge in the wave phase. In the network phase, *PD* navigation and elongation-induced nematic ordering together produce the visible network; in the wave phase, the same components continue to operate, now under the additional constraint of a food gradient that directs collective movement forward. In the model, the cryptic network during the wave phase is therefore not a memory or preservation of the network phase, but a fresh production of the same spatial process in a changed environmental context. When the wave starts from a structureless front on the food patch without a preceding visible network, a cryptic network still emerges inside the wave (Supplementary Fig. 4), so the two collective phases share the generative program rather than the pattern it produces. Removing food chemotaxis from the model recovers continuously remodeling visible networks without a traveling wave (Fig. 3e), closely resembling the dynamic network states described by Sugi et al.^12^, so within the model food chemotaxis is what converts density-driven visible network dynamics into the traveling wave.

In this sense, the relationship we observe between the wave and its cryptic network is closer in character to mycorrhizal fungi systems where the advancing hyphal front and the network it constructs are related aspects of the same ongoing process^20^, than to systems in which one hidden pattern shapes the dynamics of another, such as the coherent structures that organize transport in fluid flows^23,24^ or the interaction networks through which waves of behavioral change spread in fish schools^25^. Although our wave phase showed a cryptic network across a broad range of experimental conditions (Supplementary Fig. 2), no network has been reported in swarms of *C. elegans* studied at very high population density or with altered neuromodulation^8,14,15^. Whether a cryptic network is present in those swarms remains an open question.

In the wurmuration model, all four components are present in both network and wave phases, and as the food gradient changes, the same program produces a visible network off food and a cryptic network on it. This finding has implications beyond *C. elegans*: where distinct collective phases are governed by a single set of generative components, the spatial organization of one phase may be cryptically present in another, invisible to direct observation but recoverable through the right measurement perspective. Collective spatial organization may therefore sometimes be more informatively described at the level of the generative components than at the level of the patterns they produce.

## Methods

### Behavior experiment and video acquisition

#### Experimental Procedures

*C. elegans* strains N2 (laboratory reference strain) and DA609 [*npr-1(ad609)*X] were cultured at 20°C on nematode growth medium (NGM) plates seeded with *E. coli* OP50. Synchronized Day 1 adult populations were obtained by alkaline-hypochlorite treatment of gravid adults to isolate eggs, and letting eggs hatch overnight in M9 at 20°C on a rotator. L1 diapause larvae were refed on OP50 plates for 65 hours to reach adulthood. Imaging plates were prepared by pouring 100 mL of no-peptone NGM agar into 150 mm-diameter glass Petri dishes (Rotilabo KKA1.1, Carl Roth), and drying overnight on the bench. Rectangular food lawns (∼110 × 30 mm) were created by spreading 3 mL of OP50 (OD₆₀₀ ≈ 2.0) with a glass spreader. Seeded imaging plates were allowed to dry for 2-3 h at room temperature and incubated overnight at 37°C, before returning to room temperature prior to the experiment. For each experiment, ∼20,000 synchronized Day 1 adults were harvested from culture plates and washed in M9 to remove bacteria. After the final wash, worms were concentrated to a ∼200 µL pellet by centrifugation (Eppendorf Centrifuge 5702) at 2,000 rpm for 2 min, and deposited at the left edge of the food lawn using a glass pipette. After letting the worm pellet dry for 10-15 min, plates were transferred to an Axio Zoom V.16 microscope (Zeiss) for behavioral imaging.

#### Video Acquisition

Bright-field imaging was performed at 20°C using an Axio Zoom V.16 microscope (Zeiss) equipped with an Axiocam 712 color camera (Zeiss) and an HXP 200C light source (Zeiss). Imaging with a PlanApo Z 0.5x objective at 3.5x magnification gives a field of view of 68 × 51 mm. Time-lapse videos were acquired at 0.1 fps with 0.5 ms exposure at 70% light intensity using ZEN 3.4 Pro software (Zeiss). The main dataset comprises all N2 (*n*=14) and *npr-1* (*n*=8) experiments that followed this protocol, which were independently conducted on multiple days. Experiments with variations of this protocol in strain genetics, population composition, arena size or food conditions (Supplementary Fig. 2), are not pooled with the main dataset. A network phase was observed in 19 out of 22 total experiments; in the remaining three (two N2, one *npr-1*), recording began too late to show whether a network phase had formed. A wave phase formed in all 22 experiments.

#### No-food recordings

A separate set of recordings was acquired without food to document the network phase in isolation. The experimental protocol was identical to the above except that the plates were unseeded and contained 100 mL or 200 mL of no-peptone NGM agar; imaging used a single field of view centered on the worm deposition site and lasted approximately 1 h per recording. Three independent experiments were conducted for each combination of strain (N2 and *npr-1*) and agar volume. Visible network formation was observed in all twelve recordings, and no wave formed in any of them; the stability and the extent of the visible networks depended on the experimental conditions. These recordings are not pooled with the main dataset and are shown as representative snapshots in Supplementary Fig. 6.

### Video Processing and Analysis

#### Wave Metric Extraction

Wave temporal boundaries were identified from the mode first-visit frame index at the lateral extremes of the food patch, refined by manual annotation where needed. Wave masks were extracted at 10-minute intervals by morphological processing of binarized frames; centerlines were computed as row-wise averages of mask pixels, smoothed by moving average. *Wave speed* is the distance between the wave’s start and end positions divided by the time between them. These positions were annotated manually in the experiments to avoid artifacts caused by irregularities in the wave shape, and detected automatically in the simulations. *Wave bulkiness* is the maximal inscribed radius of the wave mask, averaged over the sampled frames. Additionally, to verify independently that the wave meets the two defining features of a traveling wave, the front was located automatically in each sampled frame as the leading half-maximum crossing of the density profile along the direction of advance, smoothed over ∼1 mm, and front speed was taken as the slope of a linear fit of leading-edge position against time (Supplementary Fig. 1; Supplementary Methods, Front kinematics). Frames in which the detected leading edge lay more than 2 mm behind the furthest position it had already reached were treated as detection failures and excluded from the fit; visual inspection confirmed that the worms did not move back in these frames. Full details are in Supplementary Methods, Wave Metric Extraction.

#### Wave phase duration

Wave phase durations, measured between the wave temporal boundaries defined above, ranged from 90 to 335 min and were statistically indistinguishable between strains (N2: 161 ± 56 min; *npr-1*: 193 ± 76 min; *P* = 0.290, Mann–Whitney). *Wave bulkiness*, *large hole area* and *TCI bulkiness* showed no consistent monotonic dependence on wave phase duration (Supplementary Fig. 7; Supplementary Methods, Independence of the metrics from wave phase duration). *Wave speed* is shown for completeness only, because the wave phase runs from where the wave is first seen to where it is last seen, so a slower wave takes longer to cover that stretch and a negative correlation is expected (Supplementary Fig. 7a).

#### Coverage Geometry Extraction

The TCI for each experimental replicate is generated by stacking all video frames through temporal minimum projection: each pixel retains the darkest value it ever reached across that experiment, corresponding to its moment of peak worm visitation. To suppress transient single-worm events before projection, frames are temporally median-filtered in a sliding window (6 subsamples per 10-minute window) prior to taking the minimum (parameter choice validated in Supplementary Fig. 8; see Supplementary Methods, TCI parameter validation). Within the manually annotated food patch boundary, bright worm-free regions in the TCI were segmented as holes by Otsu thresholding; components smaller than a minimum area threshold or touching >15% of the patch border were excluded. *Large hole area* (5th-rank hole by area; rank stability validated in Supplementary Fig. 9) and *TCI bulkiness* (maximal inscribed radius) of the worm-visited mask were the coverage geometry metrics. Full pipeline details are in Supplementary Methods, Coverage Geometry Extraction.

### Agent-Based Simulation

#### Model construction

A discrete-time agent-based model was implemented on a 30 x30 mm grid (20 px/mm) with a time step of 0.25 s. The initial arena features a vertically centered, rectangular food patch (21 x 12 mm) with rounded corners on the left side, occupying the right portion of the arena, and an elliptical worm reservoir to its left, mimicking the experimental layout at a smaller scale; waves therefore advance left to right. Each agent is a chain of 20 segments (1 mm body length, 50 µm width per segment), capturing the 1:20 aspect ratio^26^; all segments contribute to local density calculations, so elongated agents add a directional footprint to the density field. Forward movement adds a new segment at the head and removes the oldest one at the tail. Agents sense the local worm density, and the food odor and the local food concentration at the head. No alignment or attraction term is implemented. Agents switch stochastically between dwelling (full immobility) and roaming (head-led movement)^27,28^ with probabilities that depend on local density relative to *PD*. Simulated bouts last at most 100 s dwelling and 50 s roaming, shorter than the minutes-scale bouts reported for isolated animals^29^ (Supplementary Methods, Numerical illustration of component interactions, Effective speed and bout durations; Supplementary Note 1). Roaming steps follow a priority-ordered hierarchy of odor guidance, density-gradient navigation, and correlated random walk, with angular noise drawn uniformly from [−45°, +45°] added in all cases. Food depletes progressively at agent heads; when multiple agents occupy the same location, each individual consumes less than it would alone. To represent N2 and *npr-1* we set *PD* = 6 and *PD* = 12 segments/pixel respectively, with all other parameters constant (Supplementary Table 3). Necessity of each component was tested by leave-one-out simulations removing body elongation, roaming-dwelling switching, *PD* navigation, or food chemotaxis in turn. Full model description is in Supplementary Methods, Agent-Based Simulation.

#### Model behavior

The following describes how body elongation, roaming-dwelling switching, *PD* navigation and food chemotaxis interact dynamically across the arena. The switching probability depends on local crowding: dwelling agents in sparse regions are likely to start roaming, roaming agents near their preferred local worm density are likely to stop and dwell, and roaming agents in overly dense regions are unlikely to stop and are pushed back into motion by overcrowding. This single mechanism spontaneously drives the population toward *PD* without any explicit attraction rule. If an agent is roaming, it then asks, in order: do I smell food and am I not already on it? With a probability that rises with odor magnitude and falls with local food concentration, it roams up the gradient, so chemotaxis is suppressed on intact food and released as food is depleted. Agents that do not enter this odor-guided mode look at the eight pixels around their head: if a density gradient exists, they step toward the pixel whose density is closest to *PD*, with over-dense pixels penalized threefold (Supplementary Methods, Agent-Based Simulation, Density-gradient navigation). If no gradient is found because the neighborhood is uniform or empty, they continue in their current direction with a nudge of noise. Between steps, the agents’ collective positions continuously reshape the local worm density field and food consumption carves depletion into the patch and thereby shifts the odor gradient that guides chemotaxis. The body geometry turns these interactions into spatial order. Elongated agents add a directional footprint to the density field, which gives rise to nematic alignment and ultimately to the visible network in the network phase and the cryptic network in the wave phase.

#### Simulation Analysis

Four types of simulation sets were generated: baseline comparison sets (N2 and *npr-1*, 20 replicates each; Fig. 4), leave-one-out sets (each component removed in turn, shown as examples in Fig. 3), a sensitivity analysis (±20% perturbations, one parameter at a time, 40 replicates per condition; Supplementary Fig. 5; Supplementary Methods, Sensitivity analysis), and a no-network-phase control (no off-food reservoir; the wave starts from a structureless front made of agents seeded randomly in a narrow strip spanning the initial 5% of the food patch; 20 replicates at each *PD*; Supplementary Fig. 4). Simulated videos, rendered at 0.1 fps to match the experimental frame rate, were processed through the same TCI and metric extraction pipeline as the experimental data, with two settings that depend on spatial resolution: the pixel scale (30 px/mm for experiments, 20 px/mm for simulations) and the minimum hole area (100 px for experiments and 10 px for simulations). All metrics are reported in physical units (Supplementary Methods, Coverage Geometry Extraction). The close-up views in Fig. 2b were rendered separately from a 6 x 6 mm window inside the food patch, recorded from the moment the wave entered it and sampling every tenth simulation step (2.5 s), so that individual agents and immobile dwelling agents can be seen. Nematic alignment was quantified from cross section-corrected angular distributions before and after pairwise interactions (Supplementary Methods, Agent-Based Simulation, Nematic alignment analysis). For comparisons to experimental data in Fig. 4a,c, metric values within each experimental dataset or simulation set were independently min–max normalized to [0–1] over the combined N2 and *npr-1* range; this normalization does not affect the reported *P* values (see Statistics and reproducibility). Full statistical analysis details are in Supplementary Tables 2 and 4.

### Statistics and reproducibility

Analyses were performed in MATLAB R2025b (MathWorks) with the Image Processing and Statistics and Machine Learning toolboxes and the CircStat toolbox^30^. The wave metrics (*wave speed*, *wave bulkiness*) and the coverage geometry metrics (*large hole area*, *TCI bulkiness*) were each compared between strains within the experiments and within the simulations, by distribution-free permutation tests on the difference in means (10,000 permutations). Differences in variability between replicates were tested in the same way on the log ratio of standard deviations (100,000 permutations). Effect sizes are Hedges’ *g*. Correlations are Spearman’s ρ; nematic alignment is summarized by the mean resultant length of doubled angles, with the Rayleigh test. All comparisons and correlations are two-sided. *P* values of these eight comparisons are Bonferroni-corrected within pairs, one for each metric type (wave metrics, coverage geometry) in the experiments and in the simulations. The min–max normalization in Fig. 4a,c is for display only and leaves *P* values and effect sizes unchanged. Values are mean ± s.d., and *P* values below 0.001 are reported as *P* < 0.001.

Unless stated otherwise, *n* is the number of independent experiments or independent simulation runs. No statistical method was used to predetermine sample size. Groups were defined by strain, and the investigators knew the strain identity of each recording during acquisition and annotation. Metric extraction is automated and uses identical parameters for both strains. Experimental *wave speed* is the only metric computed from manually annotated start and end positions, and it agrees closely with an automatic measure, the front speed from kymographs of the raw videos (Results; Supplementary Fig. 1). All statistics were generated in one seeded run of the deposited analysis script (Code availability); the full table of statistics is given in Supplementary Data 1.

## Supporting information

Full table of statistics

## AI Disclosure

The authors used large language model tools (Claude, ChatGPT and Gemini) as assistants throughout coding, analysis and writing, in an iterative process under their direction. They designed the study and the model, checked all code, text and reported values, and take full responsibility for the content of this article.

## Data Availability

Datasets used in this study will be available at Zenodo upon publication (https://doi.org/10.5281/zenodo.20506817), including the compressed videos and TCIs of the main dataset, the TCIs of Supplementary Fig. 2, and example simulation videos and TCIs. The full resolution raw data are available from the corresponding author on request. Source data containing numerical values underlying the figures are provided with this paper.

## Code Availability

All custom MATLAB codes used in this study are available at https://github.com/SerenaDingLab/Wurmuration_2026, including the agent-based simulation, TCI generation pipeline, wave and coverage geometry metric extraction, and figure generation scripts.

## Acknowledgments

We thank Mark Blaxter for naming wurmuration, and Patricia Mütze for performing manual wave annotations and for assistance with an initial figure preparation draft. We thank Linus Schumacher, Harikrishnan Rajendran and Antonio Costa for critical reading of the manuscript. Some strains were provided by the CGC, which is funded by NIH Office of Research Infrastructure Programs (P40 OD010440). This study was funded through the Max Planck Institute of Animal Behavior, the Deutsche Forschungsgemeinschaft (DFG, German Research Foundation) under Germany’s Excellence Strategy – EXC 2117 – 422037984, and the BABOTs consortium grant (Horizon Europe, European Innovation Council Pathfinder Work Programme) under grant agreement no. 101098722.

## Author Contributions

A.P.: Conceptualization, Methodology, Data curation, Formal analysis, Software, Visualization, Writing - original draft. S.S.: Investigation, Methodology, Data curation, Writing - review & editing. G.R.: Investigation. S.S.D.: Conceptualization, Funding acquisition, Methodology, Project administration, Resources, Supervision, Writing - review & editing.

## References

1. Vicsek, T., Czirók, A., Ben-Jacob, E., Cohen, I. & Shochet, O. Novel Type of Phase Transition in a System of Self-Driven Particles. Phys. Rev. Lett. 75, 1226–1229 (1995).

2. Vicsek, T. & Zafeiris, A. Collective motion. Phys. Rep. 517, 71–140 (2012).

3. Sumpter, D. J. T. The principles of collective animal behaviour. Philos. Trans. R. Soc. B Biol. Sci. 361, 5–22 (2005).

4. Gregor, T., Fujimoto, K., Masaki, N. & Sawai, S. The onset of collective behavior in social amoebae. Science 328, 1021–1025 (2010).

5. Buhl, C. et al. From Disorder to Order in Marching Locusts. Science 312, 1402–1406 (2006).

6. Tunstrøm, K. et al. Collective States, Multistability and Transitional Behavior in Schooling Fish. PLOS Comput. Biol. 9, e1002915 (2013).

7. Couzin, I. D., Krause, J., James, R., GD, R. & Franks, N. R. Collective memory and spatial sorting in animal groups. J Theor Biol 218, 1–11 (2002).

8. Demir, E., Yaman, Y. I., Kocabas, A. & Basaran, M. Dynamics of pattern formation and emergence of swarming in Caenorhabditis elegans. eLife 9, 52781 (2020).

9. Ding, S. S., Schumacher, L. J., Javer, A. E., Endres, R. G. & Brown, A. E. Shared behavioral mechanisms underlie C. elegans aggregation and swarming. eLife 8, e43318 (2019).

10. Gray, J. M. et al. Oxygen sensation and social feeding mediated by a C. elegans guanylate cyclase homologue. Nature 430, 317–322 (2004).

11. Rogers, C., Persson, A., Cheung, B. & de Bono, M. Behavioral Motifs and Neural Pathways Coordinating O2 Responses and Aggregation in C. elegans. Curr. Biol. 16, 649– 659 (2006).

12. Sugi, T., Ito, H., Nishimura, M. & Nagai, K. H. C. elegans collectively forms dynamical networks. Nat. Commun. 10, 683 (2019).

13. Bono, M. de & Bargmann, C. I. Natural Variation in a Neuropeptide Y Receptor Homolog Modifies Social Behavior and Food Response in C. elegans. Cell 94, 679–689 (1998).

14. Ding, S. S., Romenskyy, M., Sarkisyan, K. S. & Brown, A. E. X. Measuring Caenorhabditis elegans Spatial Foraging and Food Intake Using Bioluminescent Bacteria. Genetics 214, 577–587 (2020).

15. Shahi, N. et al. Neuromodulation of swarming behavior in Caenorhabditis elegans: Insights into the conserved role of calsyntenins. Proc. Natl. Acad. Sci. 123, e2520029123 (2026).

16. Keller, E. F. & Segel, L. A. Traveling bands of chemotactic bacteria: A theoretical analysis. J. Theor. Biol. 30, 235–248 (1971).

17. Adler, J. Chemotaxis in Bacteria. Science 153, 708–716 (1966).

18. Cates, M. E. & Tailleur, J. Motility-Induced Phase Separation. Annu. Rev. Condens. Matter Phys. 6, 219–244 (2015).

19. Marchetti, M. C. et al. Hydrodynamics of soft active matter. Rev. Mod. Phys. 85, 1143– 1189 (2013).

20. Oyarte Galvez, L., et al. A travelling-wave strategy for plant–fungal trade. Nature 639, 172–180 (2025).

21. Coates, J. C. & de Bono, M. Antagonistic pathways in neurons exposed to body fluid regulate social feeding in Caenorhabditis elegans. Nature 419, 925–929 (2002).

22. Chen, Y. & Ferrell, J. E. C. elegans colony formation as a condensation phenomenon. Nat. Commun. 12, 4947 (2021).

23. Haller, G. & Yuan, G. Lagrangian coherent structures and mixing in two-dimensional turbulence. Phys. Nonlinear Phenom. 147, 352–370 (2000).

24. Peacock, T. & Haller, G. Lagrangian coherent structures: The hidden skeleton of fluid flows. Phys. Today 66, 41–47 (2013).

25. Rosenthal, S. B., Twomey, C. R., Hartnett, A. T., Wu, H. S. & Couzin, I. D. Revealing the hidden networks of interaction in mobile animal groups allows prediction of complex behavioral contagion. Proc. Natl. Acad. Sci. 112, 4690–4695 (2015).

26. Brenner, S. The genetics of Caenorhabditis elegans. Genetics 77, 71–94 (1974).

27. Fujiwara, M., Sengupta, P. & McIntire, S. L. Regulation of Body Size and Behavioral State of C. elegans by Sensory Perception and the EGL-4 cGMP-Dependent Protein Kinase. Neuron 36, 1091–1102 (2002).

28. Arous, J. B., Laffont, S. & Chatenay, D. Molecular and Sensory Basis of a Food Related Two-State Behavior in C. elegans. PLOS ONE 4, e7584 (2009).

29. Flavell, S. W. et al. Serotonin and the Neuropeptide PDF Initiate and Extend Opposing Behavioral States in C. elegans. Cell 154, 1023–1035 (2013).

30. Berens, P. CircStat: A MATLAB Toolbox for Circular Statistics. J. Stat. Softw. 31, 1–21 (2009).

