## Supplementary Information for "From Networks to Traveling Waves and Back: Re-emergent Patterns Across Collective Phases in *C. elegans*"

**Supplementary Figures** 2
Supplementary Figure 1. The wave is a single front over time of constant speed and near-stationary profile 2
Supplementary Figure 2. Cryptic network appears robustly across diverse experimental conditions 3
Supplementary Figure 3. Model architecture schematic 4
Supplementary Figure 4. The cryptic network is generated within the wave phase rather than inherited from the network phase 5
Supplementary Figure 5. The model is robust to ±20% parameter perturbations 6
Supplementary Figure 6. Populations deposited without a food patch show visible network formation but no wave 7
Supplementary Figure 7. Independence of the metrics from wave phase duration. 8
Supplementary Figure 8. TCI pipeline sensitivity 9
Supplementary Figure 9. Hole-rank metric stability 10
**Supplementary Tables** 11
Supplementary Table 1. Responses of the wave and coverage geometry metrics to ±20% perturbations 11
Supplementary Table 2. Statistical tests 12
Supplementary Table 3. Model parameters and simulation settings 13
Supplementary Table 4. Spearman correlation tests for wave metrics and coverage geometry 14
**Supplementary Methods** 15
1. Wave Metric Extraction 15
1.1 Front kinematics 15
2. Coverage Geometry Extraction 16
2.1 TCI parameter validation 16
2.2 Hole-rank metric stability 17
3. Agent-Based Simulation 18
4. Numerical illustration of component interactions 19
5. Leave-one-out simulations 21
6. Sensitivity analysis 22
7. Independence of the metrics from wave phase duration 22
**Supplementary Notes** 23
Supplementary Note 1. Biological realism of model parameters and acknowledgment of quantitative limitations 23
Supplementary Note 2. Open questions and directions for future investigation 24
**Supplementary References** 24

### Supplementary Figures

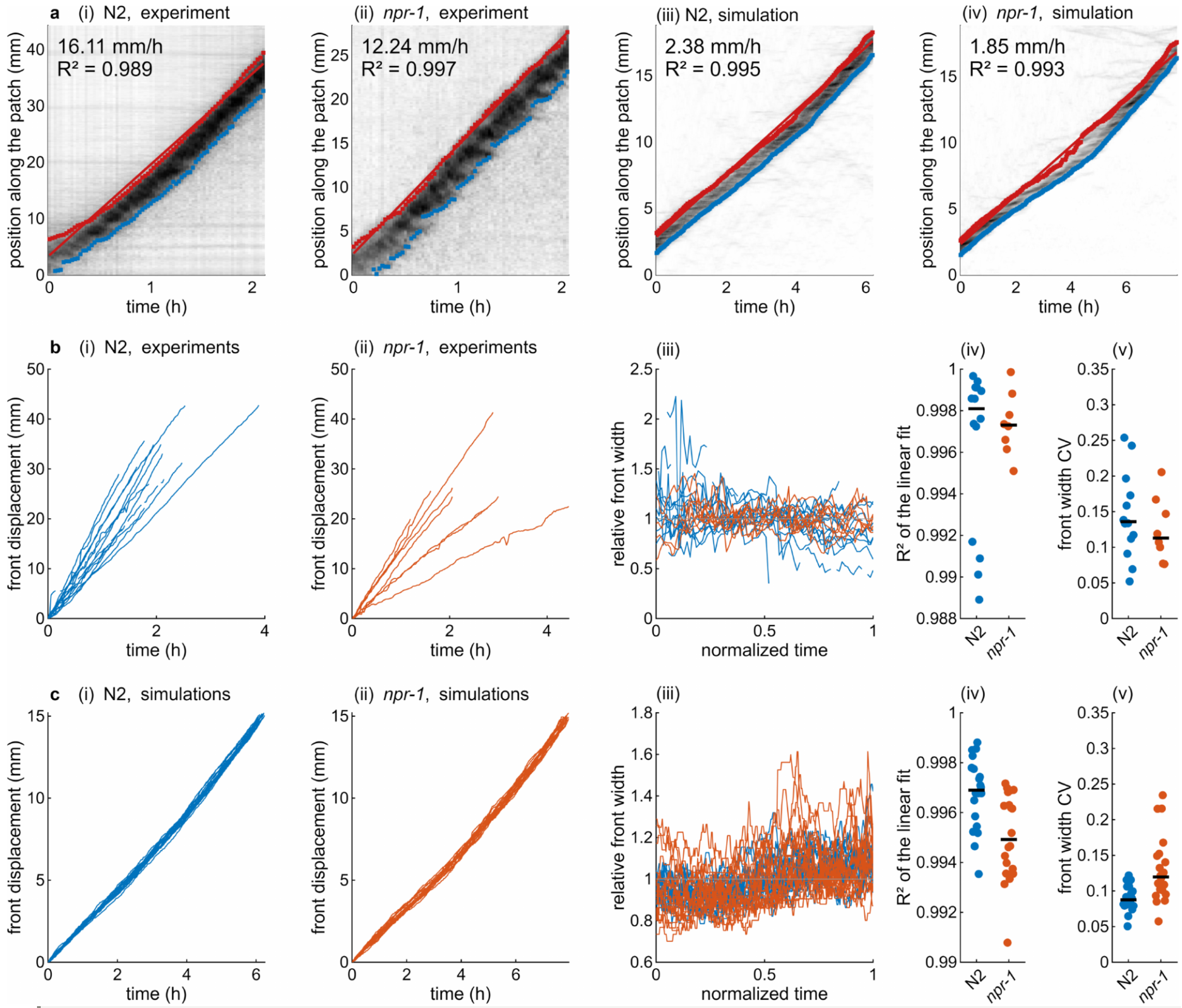


**Supplementary Figure 1. The wave is a single front over time of constant speed and near-stationary profile.** Front position and width are measured directly from the raw image series, independently of the wave metrics, as the half-maximum crossings of the density profile along the direction of advance (Supplementary Methods, Front kinematics). **a** Kymographs of one representative experimental or simulation replicate per strain, with the detected leading edge (red points), its linear fit (red line; front speed and $R^2$ in each panel) and the trailing edge (blue points): (i) N2 experiment, (ii) *npr-1* experiment, (iii) N2 simulation, (iv) *npr-1* simulation. **b** Experiments: front displacement against time for all replicates, (i) N2 and (ii) *npr-1*; (iii) front width relative to each replicate's mean, against normalized time (N2 blue, *npr-1* orange); (iv) $R^2$ of the linear fit and (v) coefficient of variation (CV) of front width, per replicate, with medians (black lines). **c** The same for the simulations. The front advances linearly in every experimental and simulation replicate (median $R^2$ 0.995–0.998) and both its edges are detected in a median 93% (N2) and 90% (*npr-1*) of experimental frames and in every simulated frame. Front width varies little within a replicate (median CV 0.09–0.14), apart from a slow drift in three of the four groups (Supplementary Methods, Front kinematics). $n$ = 14 N2 and 8 *npr-1* experimental replicates; $n$ = 20 simulation replicates per strain. Source data are provided as a Source Data file.

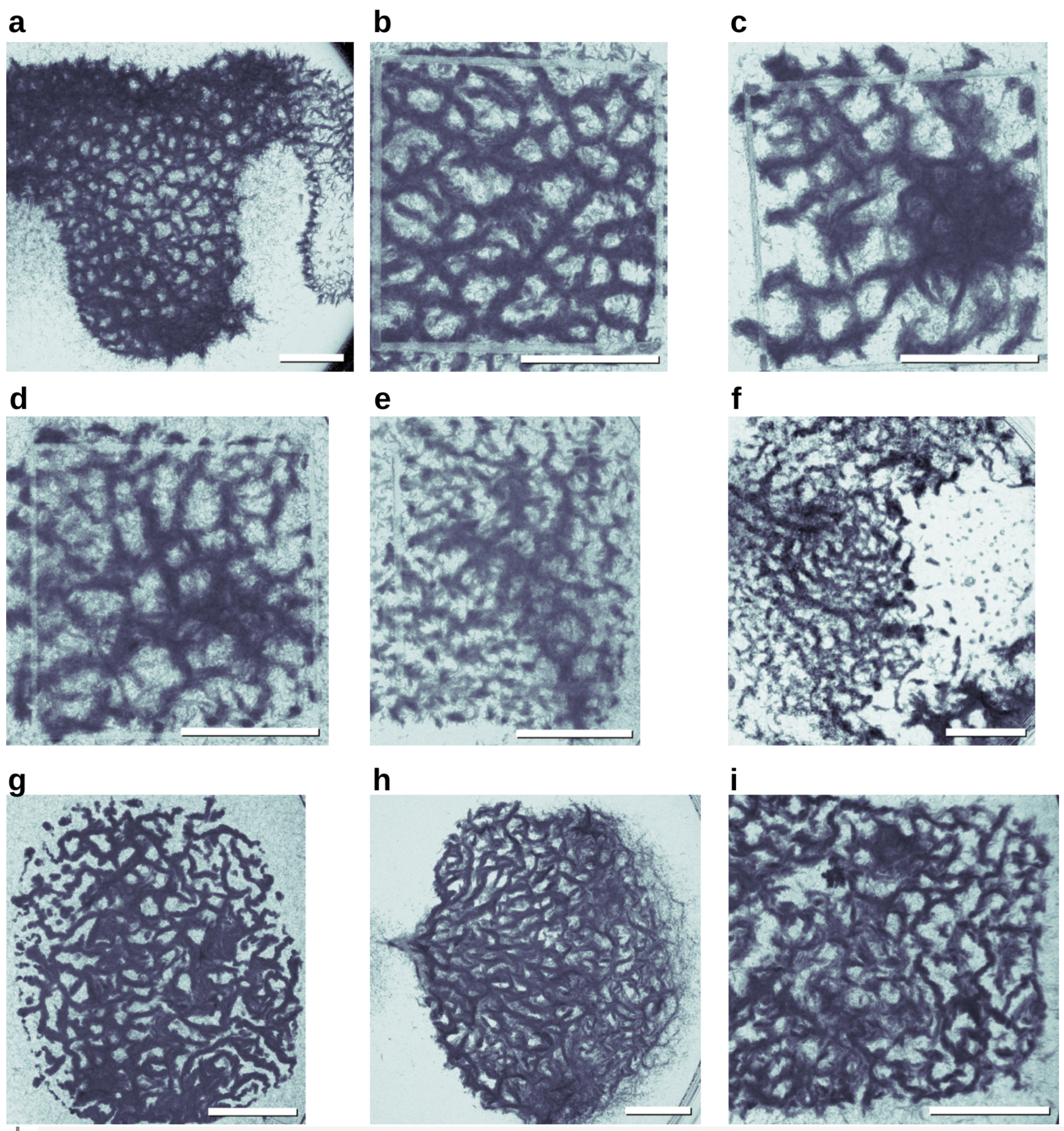


**Supplementary Figure 2. Cryptic network appears robustly across diverse experimental conditions.** All panels show TCIs generated by the same minimum-projection pipeline used for the main dataset (see Methods, Coverage Geometry Extraction). Dark bundles mark regions worms visited; bright holes are enclosed regions they left unvisited. Each panel comes from an independent experiment outside the main N2 and *npr-1* dataset, demonstrating that the cryptic network is not specific to particular strains or experimental configurations. **a** N2, 20,000 Day 1 adults, 150 mm diameter plate, snake-shaped OP50 lawn. **b** DR47 [*daf-11(m47)V*], 10,000 Day 1 adults, 55 mm diameter plate, ~20x20 mm square OP50 lawn. **c** JU1249 (wild isolate), 10,000 Day 1 adults, 55 mm diameter plate, ~20x20 mm square OP50 lawn. **d** N2, mixed-stage population 55 mm diameter plate, ~20x20 mm square OP50 lawn. **e** N2, starved population 55 mm diameter plate, ~20x20 mm square OP50 lawn. **f** OMG2 [*mIs12[myo-2p::GFP+pes-10p::GFP+F22B7.9p::GFP]II;npr-1(ad609)X*], 4,000 Day 1 adults, 55 mm diameter plate, uniform full-plate OP50 lawn. **g** DA609 [*npr-1(ad609)*X], 10,000 Day 1 adults, 90 mm diameter plate, ~35x45 mm rectangular lawn of freeze-dried OP50 (LabTIE). **h** OMG2, 12,000 Day 1 adults, 90 mm diameter plate, ~45x50 mm rectangular lawn of freeze-dried OP50 (LabTIE). **i** DA609, 10,000 Day 1 adults, 55 mm diameter plate, ~25x15 mm rectangular OP50 lawn. All scale bars: 5 mm.

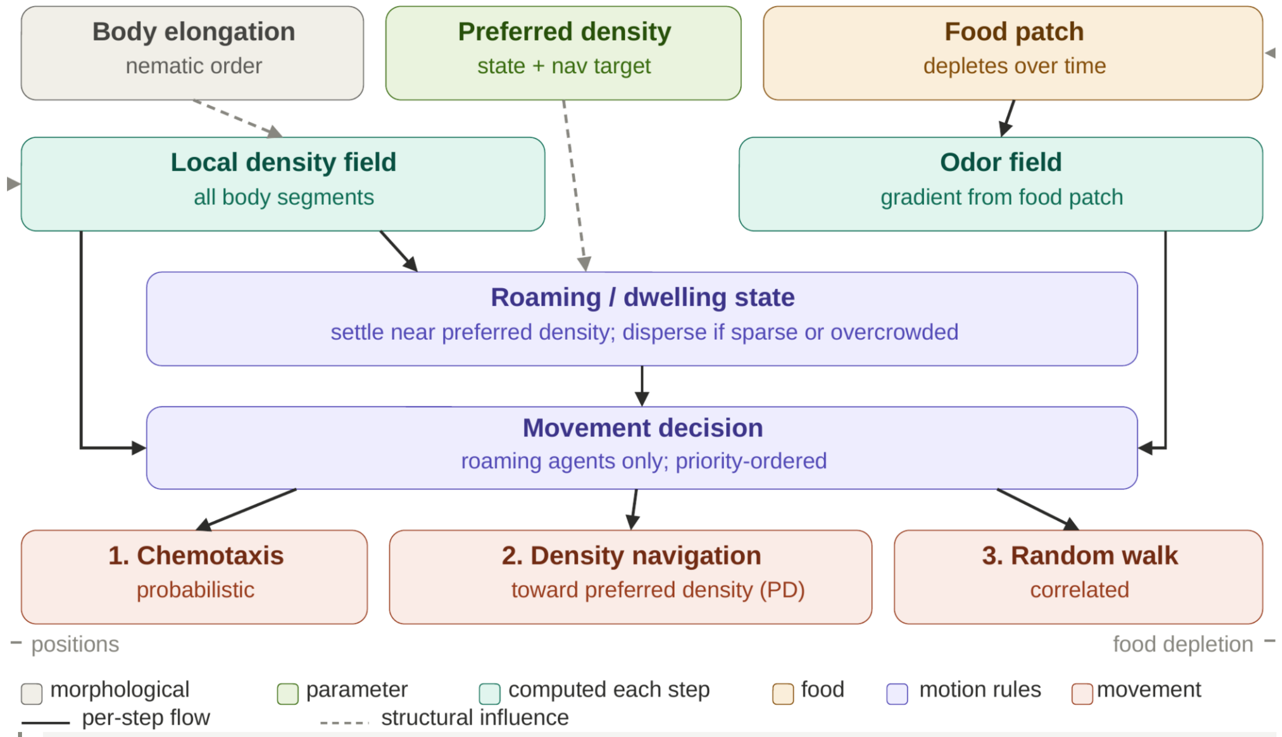


**Supplementary Figure 3. Model architecture schematic.** See Supplementary Methods, Agent-Based Simulation, for the full description.

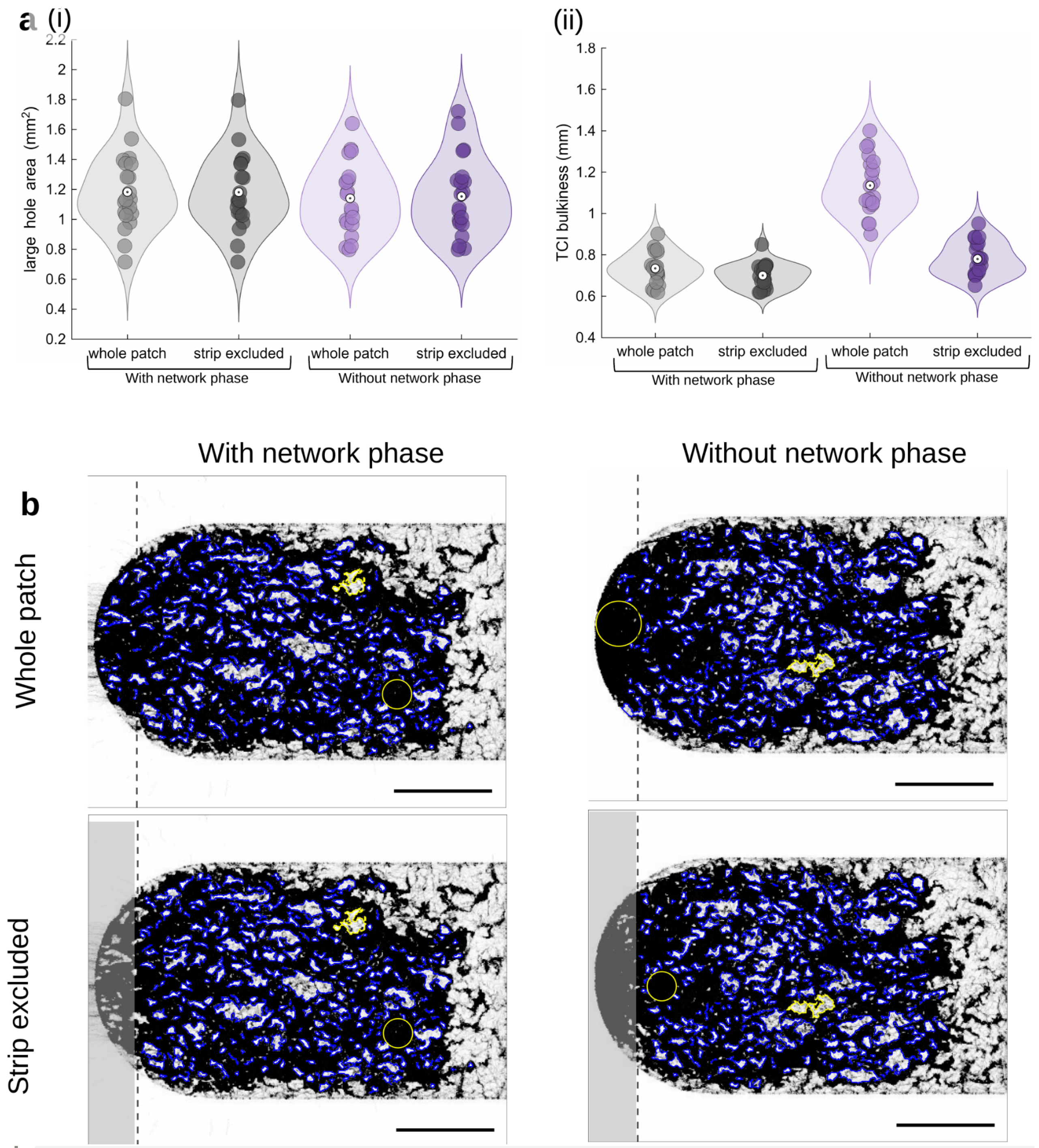


**Supplementary Figure 4. The cryptic network is generated within the wave phase rather than inherited from the network phase.** Simulations with a network phase (reservoir, network phase, then wave phase) are compared with the no-network-phase control, in which the wave starts from a structureless front made of agents seeded randomly in a narrow strip spanning the initial 5% of the food patch, with no reservoir and all else unchanged (*PD* = 12; *n* = 20 replicates each; see *PD* = 6 in Supplementary Data 1). The seeded strip appears in the TCI as a dark region that can set *TCI bulkiness*, so both conditions are also analyzed with that strip excluded. **a** *Large hole area* (i) and *TCI bulkiness* (ii), with and without a network phase, over the whole patch and with the strip exclusion; points are replicates, white circles mark the means. *Large hole area* in the control matches the value with a network phase (96–98%), and its excess *TCI bulkiness* falls from 55% to 11% once the seeded strip is excluded; percentages are ratios of condition means. **b** Representative TCIs, with network phase (left) and without network phase (right), over the whole patch (top) and with the strip exclusion (bottom; dashed line, exclusion boundary; gray areas where excluded). Blue outlines, detected holes; yellow, the fifth-largest hole and the maximal inscribed radius of the bundles. Scale bars: 5 mm. Source data are provided as a Source Data file.

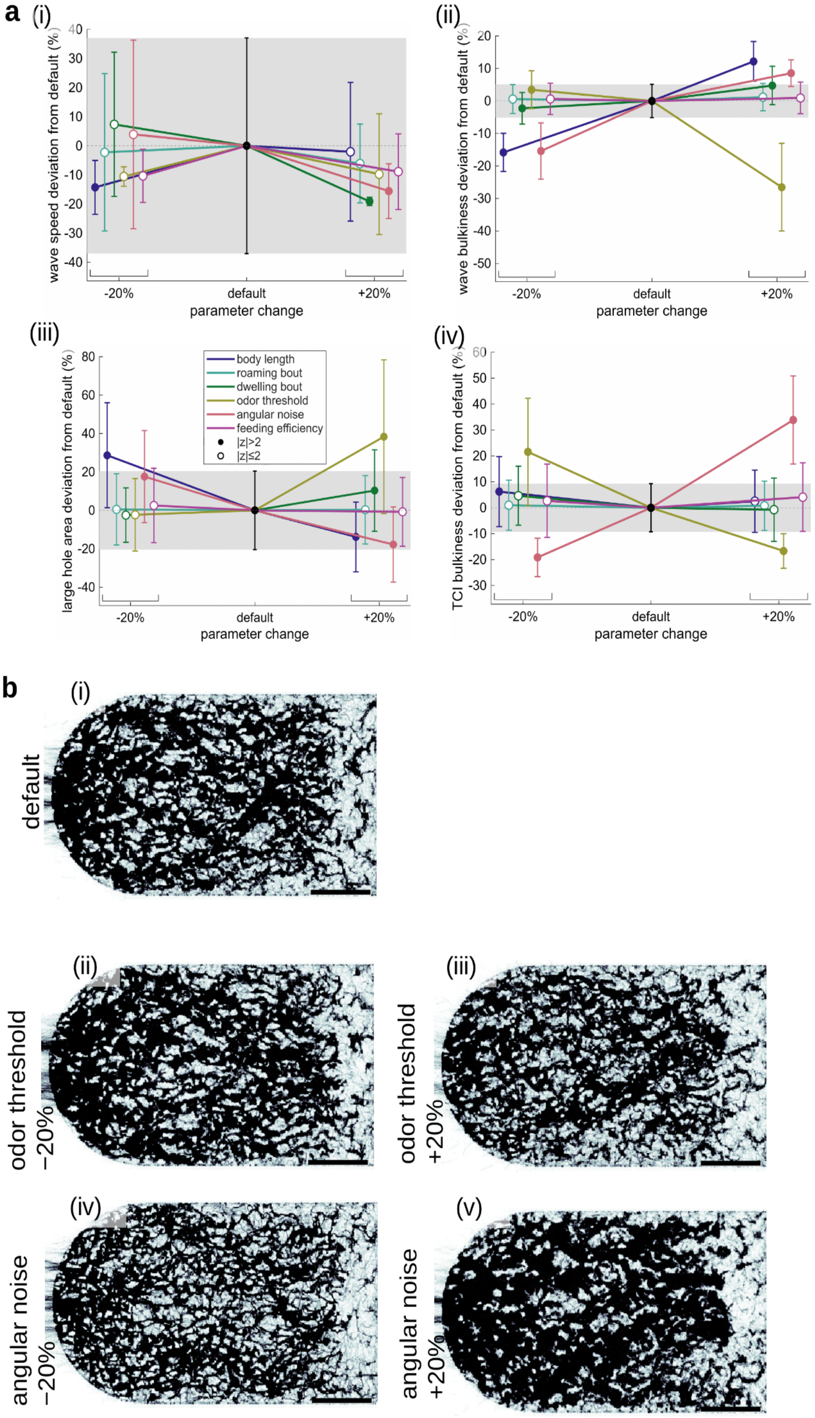


**Supplementary Figure 5. The model is robust to ±20% parameter perturbations.** Each of the parameters was varied by ±20%, one at a time, with all others at their defaults (1,500 agents, *PD* = 12; *n* = 40 replicates per condition; Supplementary Methods, Sensitivity analysis). **a** Mean deviation from the default for (i) *wave speed*, (ii) *wave bulkiness*, (iii) *large hole area* and (iv) *TCI bulkiness*. Error bars, s.d. across replicates; gray shading, s.d. across the 40 default replicates; filled markers, $|z| > 2$. Lines join each parameter's −20% and +20% responses through the default (black); markers grouped by a bracket share the same change (−20% or +20%) and are offset for visibility. **b** TCIs for the default parameterization and perturbations of the two parameters with the greatest displacement of coverage geometry (Fig. 5b), one replicate per condition: (i) default, (ii) odor threshold −20%, (iii) odor threshold +20%, (iv) angular noise −20%, (v) angular noise +20%. All scale bars: 5 mm. Source data are provided as a Source Data file.

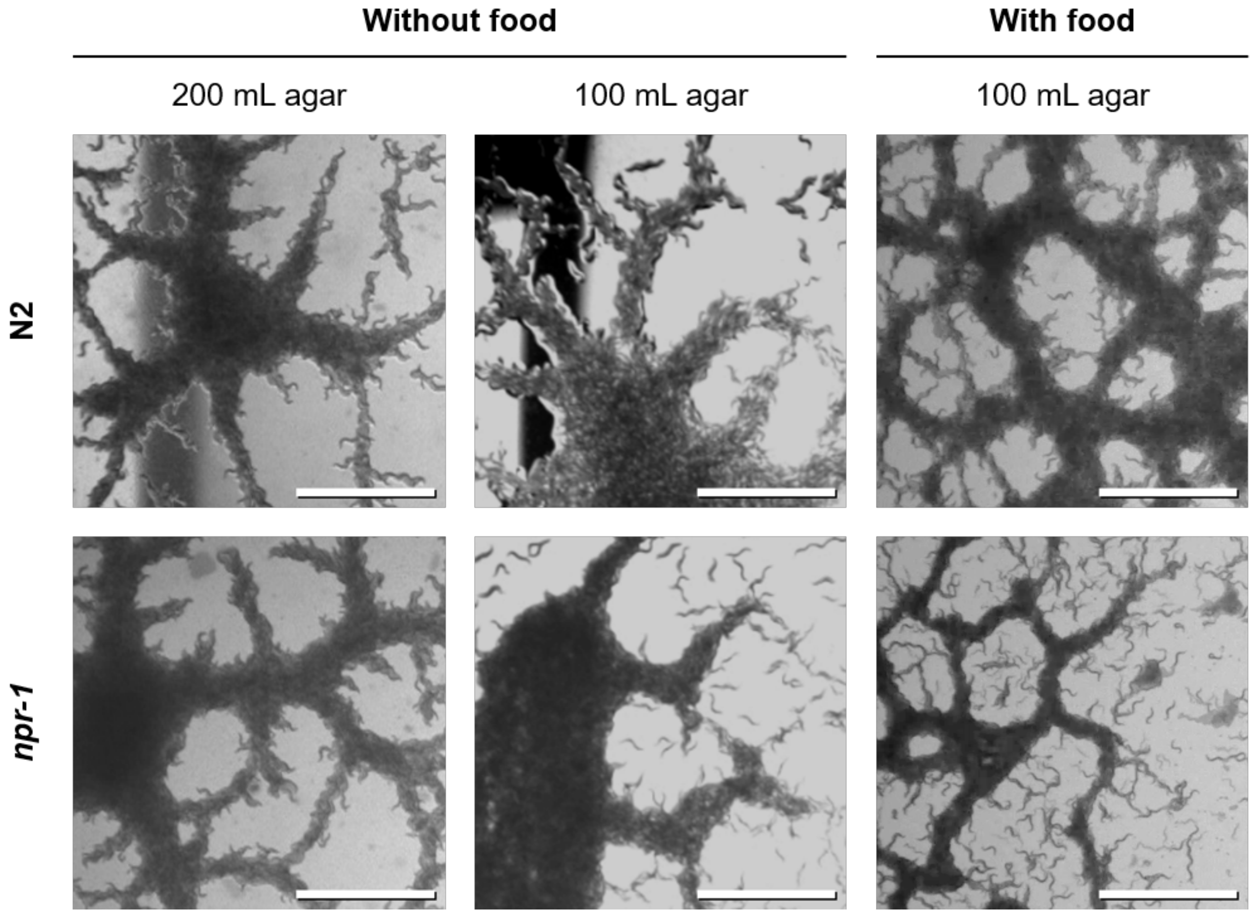


**Supplementary Figure 6. Experimental populations deposited without a food patch show visible network formation but no wave.** Sample network phase frames from independent experiments from no-food recordings (left and middle columns), or with a food patch (right column; from the main dataset). In the no-food recordings, many worms stay at the reservoir deposition site, and the visible network branches out from there. Scale bars: 5 mm.

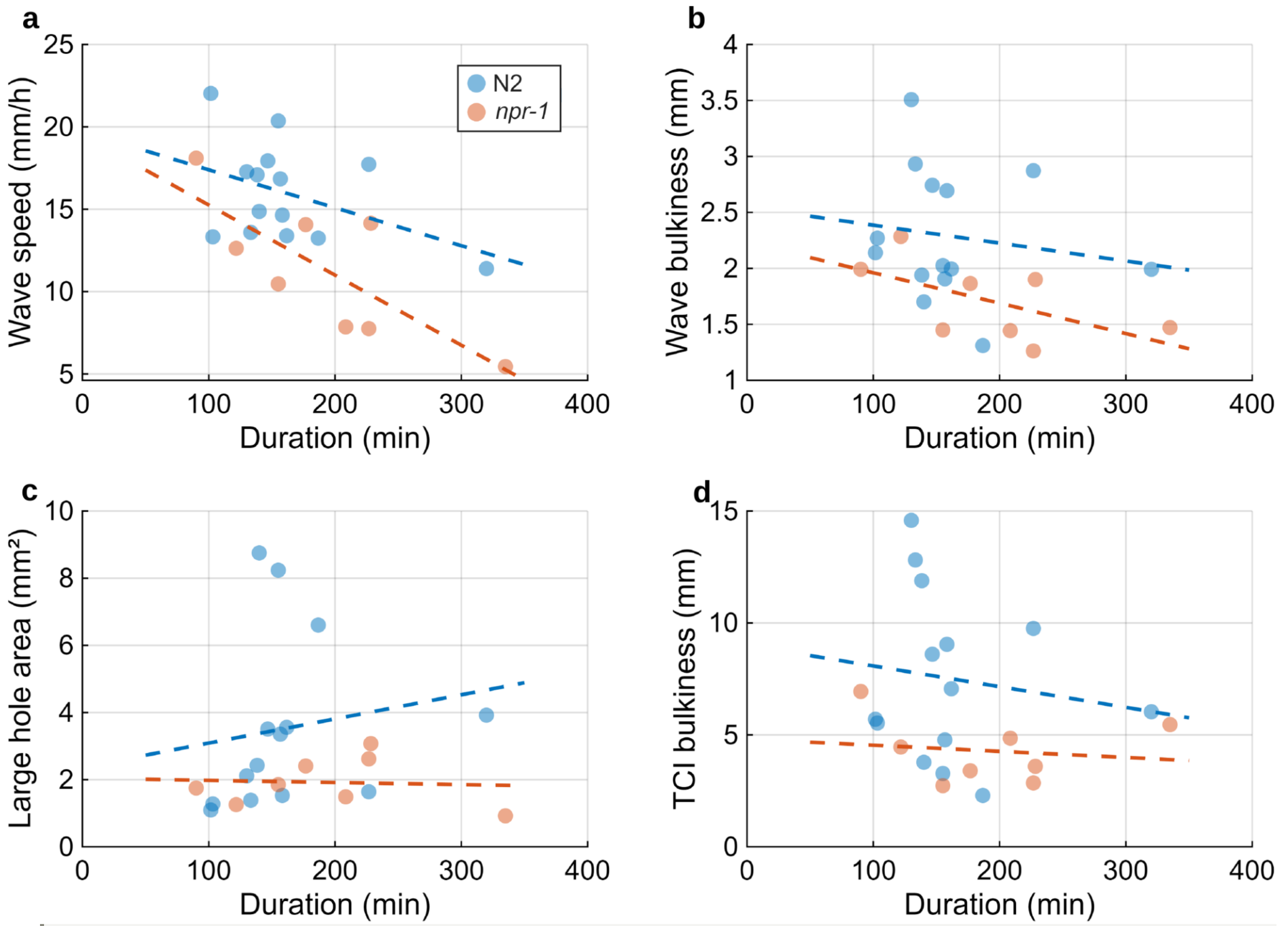


**Supplementary Figure 7. Independence of the metrics from wave phase duration.** Spearman correlations between wave phase duration and each metric, computed separately for N2 (blue, $n = 14$) and *npr-1* (orange, $n = 8$). Dashed lines are linear fits, shown for illustration; ρ and *P* values are reported per strain. **a** *Wave speed* is shown for completeness only: a slower wave takes longer to cross the patch, so a negative association is expected (N2: $\rho = -0.38$, $P = 0.19$; *npr-1*: $\rho = -0.57$, $P = 0.15$). **b** *Wave bulkiness* shows no significant correlation with wave phase duration in either strain (N2: $\rho = -0.31$, $P = 0.27$; *npr-1*: $\rho = -0.48$, $P = 0.24$). **c** *Large hole area* shows no significant correlation with wave phase duration in either strain (N2: $\rho = 0.52$, $P = 0.059$; *npr-1*: $\rho = 0.14$, $P = 0.75$). **d** *TCI bulkiness* shows no significant correlation with wave phase duration in either strain (N2: $\rho = -0.19$, $P = 0.51$; *npr-1*: $\rho = -0.07$, $P = 0.88$). Wave phase durations are statistically indistinguishable between strains (N2: 161 ± 56 min; *npr-1*: 193 ± 76 min; $P = 0.29$, Mann–Whitney), so duration does not confound the between-strain comparisons reported in the main text. Source data are provided as a Source Data file.

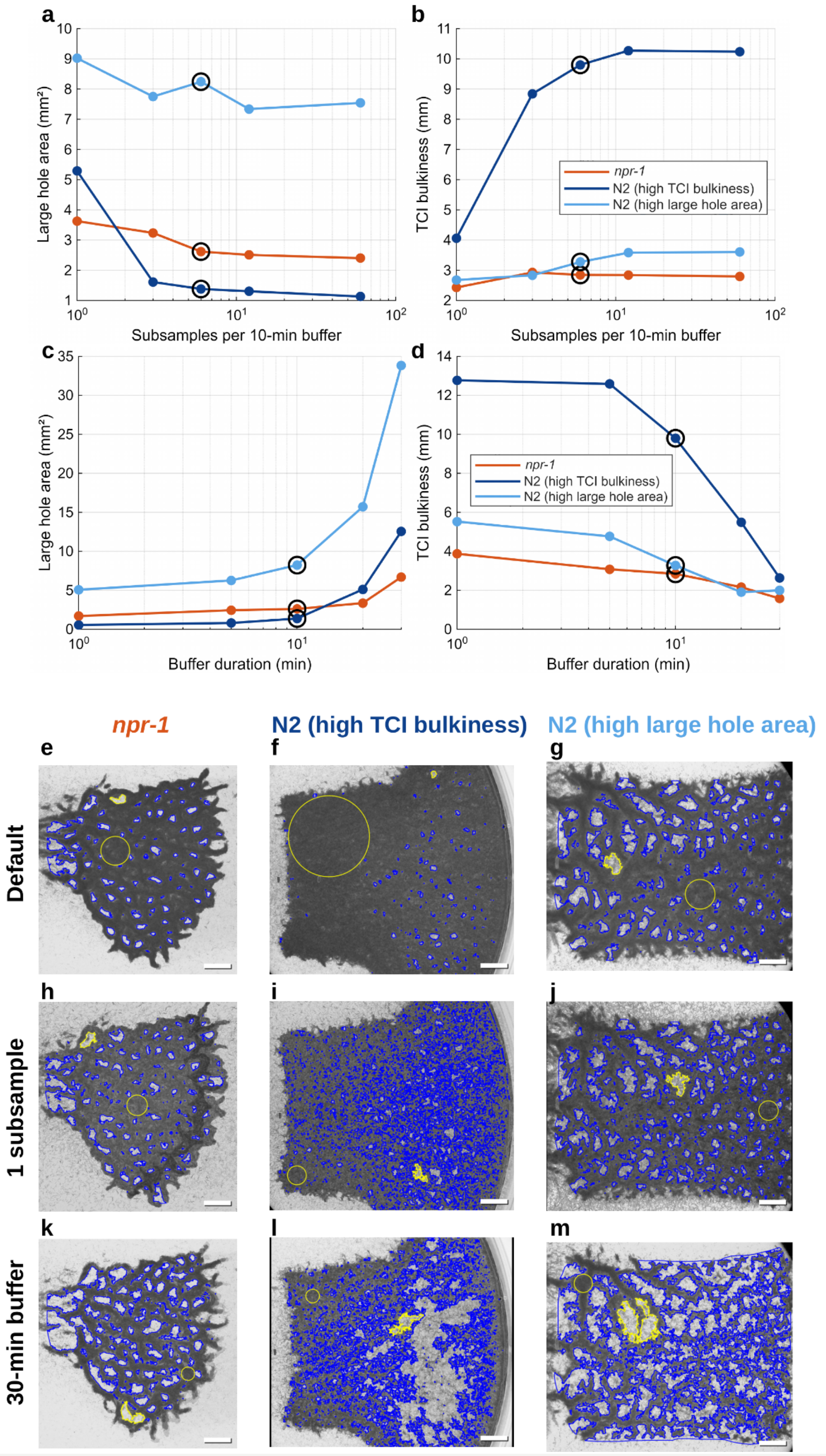


**Supplementary Figure 8. TCI pipeline sensitivity. a–d** Coverage geometry metrics (*large hole area*, *TCI bulkiness*) for three representative experiments: *npr-1* (light blue), N2 with high *TCI bulkiness* (dark blue), N2 with high *large hole area* (orange). Open black circles mark the default parameter value; x-axes in a–d are logarithmic, with swept values of 1, 3, 6, 12 and 60 subsamples in a–b and 1, 5, 10, 20 and 30 minutes in c–d. **a–b** Subsamples per 10-minute buffer, buffer duration fixed. **c–d** Buffer duration, six subsamples fixed. **e–m** TCIs with detected regions overlaid: blue outlines, all detected holes; yellow, the fifth-largest hole and the maximal inscribed radius of the bundles. All scale bars: 5 mm. Source data are provided as a Source Data file.

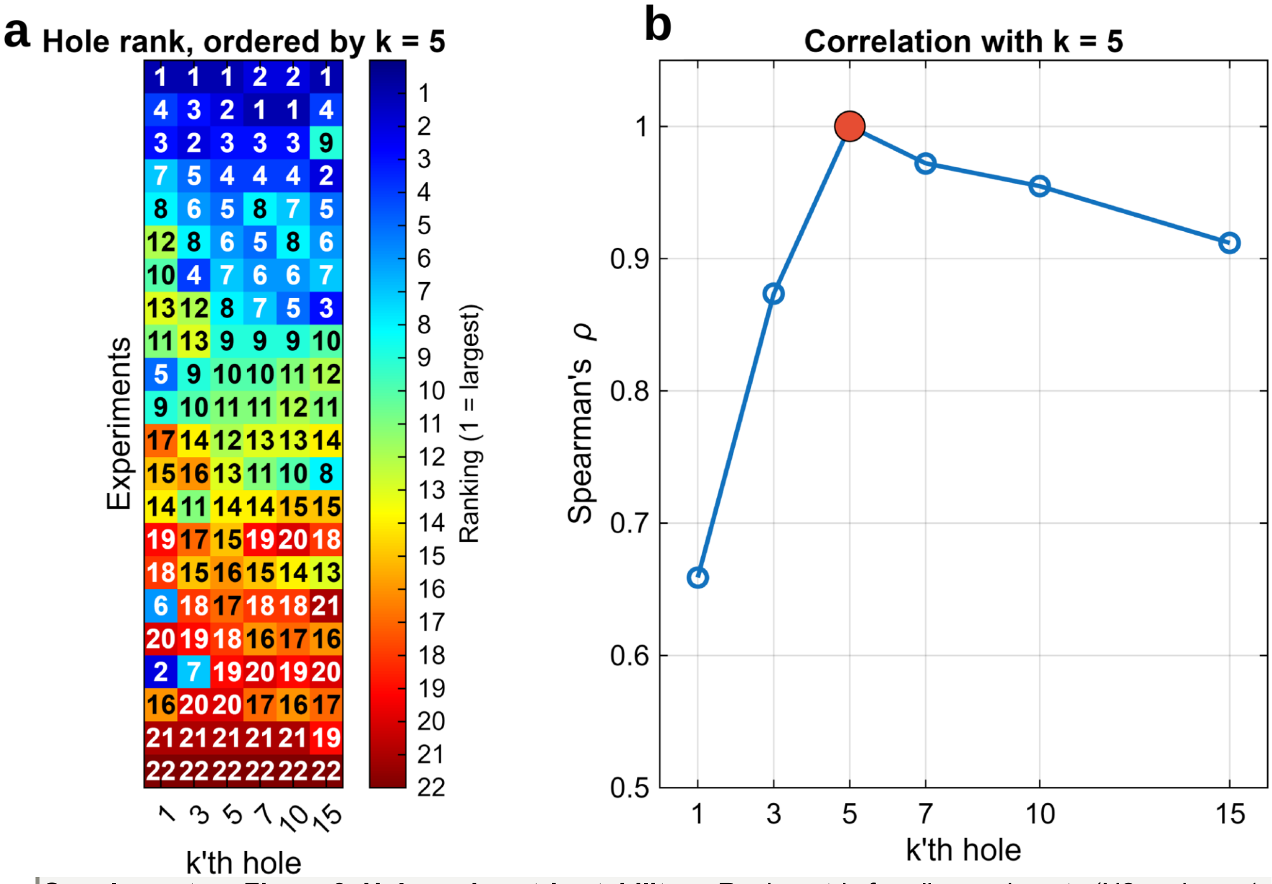


**Supplementary Figure 9. Hole-rank metric stability. a** Rank matrix for all experiments (N2 and *npr-1* combined, $n$ = 22) across hole ranks $k$ = 1, 3, 5, 7, 10, 15, ordered by the $k$ = 5 column. Color indicates rank within each column (rank 1 = largest area). **b** Spearman correlation between the $k$ = 5 ordering and each $k$. See Supplementary Methods, Coverage Geometry Extraction, Hole-rank metric stability, for interpretation. Source data are provided as a Source Data file.

### Supplementary Tables

#### Supplementary Table 1. Responses of the wave and coverage geometry metrics to ±20% perturbations

| Parameter | Change | Wave speed | Wave bulkiness | Large hole area | TCI bulkiness |
|---|---|---|---|---|---|
| Body length (segments per agent) | −20% | −14.3 ± 9.2 (−2.4)* | −15.8 ± 5.9 (−12.9)* | +28.7 ± 27.3 (+5.3)* | +6.2 ± 13.5 (+2.4)* |
| Body length (segments per agent) | +20% | −2.0 ± 23.8 (−0.3) | +12.2 ± 6.1 (+9.7)* | −13.8 ± 18.2 (−3.2)* | +2.5 ± 12.0 (+1.1) |
| Baseline roaming bout | −20% | −2.3 ± 27.0 (−0.3) | +0.6 ± 4.4 (+0.5) | +0.5 ± 18.6 (+0.1) | +1.0 ± 9.6 (+0.5) |
| Baseline roaming bout | +20% | −6.1 ± 13.5 (−1.0) | +1.2 ± 4.2 (+1.1) | +0.3 ± 17.8 (+0.1) | +0.8 ± 9.5 (+0.4) |
| Baseline dwelling bout | −20% | +7.3 ± 24.8 (+1.0) | −2.3 ± 4.9 (−2.03)* | −2.5 ± 14.2 (−0.6) | +4.6 ± 11.4 (+2.00) |
| Baseline dwelling bout | +20% | −19.1 ± 1.4 (−3.3)* | +4.8 ± 5.9 (+3.8)* | +10.3 ± 21.2 (+2.2)* | −0.7 ± 12.2 (−0.3) |
| Odor threshold | −20% | −10.6 ± 3.3 (−1.8) | +3.5 ± 5.8 (+2.8)* | −2.3 ± 18.9 (−0.5) | +21.6 ± 20.7 (+6.0)* |
| Odor threshold | +20% | −9.8 ± 20.7 (−1.5) | −26.5 ± 13.5 (−11.6)* | +38.4 ± 40.1 (+5.4)* | −16.7 ± 6.7 (−9.2)* |
| Angular noise | −20% | +3.9 ± 32.4 (+0.5) | −15.4 ± 8.7 (−9.7)* | +17.6 ± 24.0 (+3.5)* | −19.2 ± 7.4 (−10.2)* |
| Angular noise | +20% | −15.6 ± 9.4 (−2.6)* | +8.5 ± 4.1 (+8.2)* | −17.8 ± 19.6 (−4.0)* | +33.9 ± 17.0 (+11.1)* |
| Collective feeding efficiency | −20% | −10.3 ± 9.1 (−1.7) | +0.6 ± 4.8 (+0.6) | +2.5 ± 19.3 (+0.6) | +2.7 ± 14.1 (+1.0) |
| Collective feeding efficiency | +20% | −8.9 ± 13.0 (−1.4) | +0.9 ± 4.9 (+0.8) | −0.8 ± 17.9 (−0.2) | +4.1 ± 13.2 (+1.6) |

Each parameter was set to 0.8x and 1.2x its default with all others held at default (Supplementary Methods, Sensitivity analysis). Entries are the deviation of the condition mean from the default mean ± the s.d. across that condition's replicates, both as a percentage of the default mean, with *z*, the shift divided by its standard error, in parentheses. An asterisk marks $|z| > 2$. Default condition, 1,500 agents at a *PD* of 12, mean ± s.d. (s.d. as a percentage of the mean): *wave speed* 2.24 ± 0.827 mm $h^{-1}$ (37.0%); *wave bulkiness* 0.300 ± 0.015 mm (5.1%); *large hole area* 1.23 ± 0.250 $mm^2$ (20.4%); *TCI bulkiness* 0.755 ± 0.070 mm (9.3%). $n = 40$ for every condition and metric.

#### Supplementary Table 2. Statistical tests

| Figure | Metric | Unit | Group 1 | Group 2 | Test | *g* | *P* | *P* (Bonferroni) |
|---|---|---|---|---|---|---|---|---|
| Figure 3a,b | Mean resultant length | *r* | Elongated agents, *r* (corrected, doubled): 0.450 after, 0.109 before interactions | — | Rayleigh | — | < 0.001 | — |
| Figure 3a,b | Mean resultant length | *r* | Round agents, *r* (doubled): 0.155 after, 0.202 before interactions | — | Rayleigh | — | < 0.001 | — |
| Figure 4 | Wave speed | mm $h^{-1}$ | exp. N2: 16.0 ± 3.00 (*n* = 14) | exp. *npr-1*: 11.3 ± 4.20 (*n* = 8) | Permutation | 1.30 | 0.006 | 0.011 |
| Figure 4 | Wave speed | mm $h^{-1}$ | sim. N2: 2.45 ± 0.025 (*n* = 20) | sim. *npr-1*: 2.15 ± 0.502 (*n* = 20) | Permutation | 0.815 | 0.006 | 0.011 |
| Figure 4 | Wave bulkiness | mm | exp. N2: 2.29 ± 0.585 (*n* = 14) | exp. *npr-1*: 1.71 ± 0.352 (*n* = 8) | Permutation | 1.08 | 0.020 | 0.041 |
| Figure 4 | Wave bulkiness | mm | sim. N2: 0.375 ± 0.016 (*n* = 20) | sim. *npr-1*: 0.308 ± 0.016 (*n* = 20) | Permutation | 4.01 | < 0.001 | < 0.001 |
| Figure 4 | Large hole area | $mm^2$ | exp. N2: 3.53 ± 2.56 (*n* = 14) | exp. *npr-1*: 1.92 ± 0.727 (*n* = 8) | Permutation | 0.733 | 0.100 | 0.201 |
| Figure 4 | Large hole area | $mm^2$ | sim. N2: 0.482 ± 0.106 (*n* = 20) | sim. *npr-1*: 1.18 ± 0.255 (*n* = 20) | Permutation | −3.52 | < 0.001 | < 0.001 |
| Figure 4 | TCI bulkiness | mm | exp. N2: 7.51 ± 3.74 (*n* = 14) | exp. *npr-1*: 4.28 ± 1.44 (*n* = 8) | Permutation | 0.991 | 0.029 | 0.058 |
| Figure 4 | TCI bulkiness | mm | sim. N2: 0.780 ± 0.074 (*n* = 20) | sim. *npr-1*: 0.734 ± 0.076 (*n* = 20) | Permutation | 0.589 | 0.061 | 0.123 |
| Figure 5a | Standard deviation ratio, N2 over *npr-1* | ratio | exp. large hole area SD ratio (permutation): 3.53 | — | Permutation | — | 0.033 | — |
| Figure 5a | Standard deviation ratio, N2 over *npr-1* | ratio | exp. TCI bulkiness SD ratio (permutation): 2.60 | — | Permutation | — | 0.021 | — |
| Figure 5a | Standard deviation ratio, N2 over *npr-1* | ratio | sim. large hole area SD ratio (permutation): 0.415 | — | Permutation | — | < 0.001 | — |
| Figure 5a | Standard deviation ratio, N2 over *npr-1* | ratio | sim. TCI bulkiness SD ratio (permutation): 0.975 | — | Permutation | — | 0.925 | — |
| Supp. Fig. 1 | Front speed, N2 vs *npr-1* | mm $h^{-1}$ | exp. N2: 16.1 ± 3.39 (*n* = 14) | exp. *npr-1*: 11.6 ± 4.23 (*n* = 8) | Permutation | 1.16 | 0.013 | — |
| Supp. Fig. 1 | Front speed, N2 vs *npr-1* | mm $h^{-1}$ | sim. N2: 2.39 ± 0.033 (*n* = 20) | sim. *npr-1*: 1.82 ± 0.092 (*n* = 20) | Permutation | 8.05 | < 0.001 | — |
| Supp. Fig. 7 | Wave phase duration | min | exp. N2: 161 ± 55.5 (*n* = 14) | exp. *npr-1*: 193 ± 75.8 (*n* = 8) | Mann-Whitney | — | 0.290 | — |

Permutation tests use 10,000 permutations, except the comparisons of variability between replicates, which use 100,000; *P* values below 0.001 are given as < 0.001. Values are mean ± s.d. unless otherwise indicated (see individual rows for *r* and fold-ratio entries). The mean resultant length *r* is computed from doubled angles so that parallel and antiparallel pairs count alike, and is corrected for body cross-section for elongated agents only (Section 3, Nematic alignment analysis). Effect sizes are Hedges' *g*, signed so that a positive value means the first group exceeds the second. P (Bonferroni) is P corrected for multiple comparisons within each pair (Methods); a dash marks a comparison outside these pairs.

### Supplementary Table 3. Model parameters and simulation settings

| Parameter | Value | Unit | Source | Model role |
|---|---|---|---|---|
| Body length | 1 | mm | M[1] | Agent body dimensions |
| Segments per agent | 20 | — | M[1] | Aspect ratio 1:20; body geometry and density sampling along the body |
| Segment spacing | 0.05 | mm | M[1] | 1 mm ÷ 20 segments |
| Time step ($\Delta t$) | 0.25 | s | L[2] | Agent moves one segment per step at roaming speed |
| Steps per second | 4 | steps $s^{-1}$ | L[2] | ~4 segments $s^{-1}$; 0.2 mm $s^{-1}$ roaming speed |
| Angular noise | 45 | degrees | F | Half-width of the uniform random change in heading per step; sets the directional persistence of locomotion |
| Baseline dwelling bout (avgDwellingSteps) | 100 | steps | F | Reciprocal of the baseline leave rate; realized bouts vary with local density (Supplementary Methods, Effective speed and bout durations) |
| Baseline roaming bout (avgRoamingSteps) | 50 | steps | F | As above |
| Transition-probability floor | 0.25 | factor | F | Minimum leave probability in either state, as a fraction of the baseline rate |
| *PD* | 6 (N2); 12 (*npr-1*) | segments $pixel^{-1}$ | F | Fitted to qualitative wave morphology; the only parameter that differs between the two simulated strains |
| Density penalty above optimal | 3 | factor | F | Overcrowding penalized more than isolation; asymmetry constant of the mismatch function |
| Food consumption rate (*r*) | 0.999 | per step | F | Per-step retention factor, calibrated to the timescale of the wave's advance |
| Collective feeding efficiency | 0.2 | — | F | Sublinear scaling of multi-agent feeding |
| Odor decay radius | 50 | pixels | F | Approximate food gradient diffusion scale |
| Odor threshold | 0.4 | normalized | F | Minimum odor for chemotaxis activation |
| Food threshold | 0.2 | normalized | F | Local food above which chemotaxis is suppressed |
| Logistic softness | 0.05 | normalized | F | Width of both logistic factors of $p_{guided}$ |
| Arena size | 30 × 30 | mm | F | Experimental layout at smaller scale |
| Grid resolution | 20 | pixels $mm^{-1}$ | S | One pixel per segment spacing (50 µm) |
| Food patch size | 21 × 12 | mm | F | Rounded rectangle, as the experimental lawn |
| Agents per simulation | 1,500 | — | F | Population size in every simulation set |
| Render frame rate | 0.1 | fps | S | Matches the experimental frame rate |
| Termination criterion | 50 | % residual food | S | In the vertical stripe 5–10% from the right edge of the arena |
| Seeding strip width (frontWidthRate) | 0.05 | fraction of patch width | S | No-network-phase control only |
| Footprint exclusion | 0.10 | fraction of patch width | S | Twice the seeding width; applied identically to every condition in that control |

Source: M = matched to published *C. elegans* measurements; L = consistent with published ethogram or literature values; F = fitted to reproduce qualitative experimental dynamics; S = simulation or analysis setting.

#### Supplementary Table 4. Spearman correlation tests for wave metrics and coverage geometry

| Figure | Variables | *n* | Coefficient (ρ) | *P* | Notes |
|---|---|---|---|---|---|
| Figure 5a | Large hole area vs TCI bulkiness | 14 | −0.525 | 0.057 | exp. N2 within-strain scatter |
| Figure 5a | Large hole area vs TCI bulkiness | 8 | −0.619 | 0.115 | exp. *npr-1* within-strain scatter |
| Figure 5a | Large hole area vs TCI bulkiness | 20 | 0.197 | 0.404 | sim. N2 within-strain scatter |
| Figure 5a | Large hole area vs TCI bulkiness | 20 | 0.108 | 0.651 | sim. *npr-1* within-strain scatter |
| Supp. Fig. 7 | Wave speed vs wave phase duration | 14 | −0.376 | 0.186 | N2 |
| Supp. Fig. 7 | Wave speed vs wave phase duration | 8 | −0.571 | 0.151 | *npr-1* |
| Supp. Fig. 7 | Wave bulkiness vs wave phase duration | 14 | −0.314 | 0.273 | N2 |
| Supp. Fig. 7 | Wave bulkiness vs wave phase duration | 8 | −0.476 | 0.243 | *npr-1* |
| Supp. Fig. 7 | Large hole area vs wave phase duration | 14 | 0.521 | 0.059 | N2 |
| Supp. Fig. 7 | Large hole area vs wave phase duration | 8 | 0.143 | 0.752 | *npr-1* |
| Supp. Fig. 7 | TCI bulkiness vs wave phase duration | 14 | −0.191 | 0.512 | N2 |
| Supp. Fig. 7 | TCI bulkiness vs wave phase duration | 8 | −0.071 | 0.882 | *npr-1* |
| Supp. Fig. 9 | Hole area, $k$ = 1 vs $k$ = 5 | 22 | 0.659 | — | Ordering vs reference $k$ = 5 |
| Supp. Fig. 9 | Hole area, $k$ = 3 vs $k$ = 5 | 22 | 0.874 | — | Ordering vs reference $k$ = 5 |
| Supp. Fig. 9 | Hole area, $k$ = 7 vs $k$ = 5 | 22 | 0.972 | — | Ordering vs reference $k$ = 5 |
| Supp. Fig. 9 | Hole area, $k$ = 10 vs $k$ = 5 | 22 | 0.955 | — | Ordering vs reference $k$ = 5 |
| Supp. Fig. 9 | Hole area, $k$ = 15 vs $k$ = 5 | 22 | 0.912 | — | Ordering vs reference $k$ = 5 |
| Supp. Fig. 1 | Front speed (from kymographs) vs wave speed | 14 | 0.930 | < 0.001 | exp. N2; within-strain agreement between the two speed estimators |
| Supp. Fig. 1 | Front speed (from kymographs) vs wave speed | 8 | 0.905 | 0.005 | exp. *npr-1*; within-strain agreement between the two speed estimators |

Correlations are Spearman. *P* values below 0.001 are given as < 0.001; a dash in the *P* column marks a descriptive coefficient, for which no test was performed.

### Supplementary Methods

#### 1. Wave Metric Extraction

**Wave temporal boundaries.** The mode first-visit frame index was computed within the leftmost 1% and rightmost 1% of the food-patch column range of the TCI, yielding automated wave start and end frames. For experiments with complex wave dynamics, these boundaries were refined by manual annotation.

**Wave mask extraction.** At 10-minute intervals, a wave mask was extracted from the corresponding raw video frame. Sampling started after a formation delay of either 100 min from the wave start or 20% of the wave phase duration where the wave phase is shorter than 100 min. Each frame was flat-field corrected, inverted, binarized using Otsu's method, and restricted to pixels inside the eroded food-patch boundary. Morphological cleaning used erosion and dilation with a disk structuring element (radius ~7 pixels at experimental imaging resolution, scaled proportionally for simulated data) to remove isolated noise and fill gaps. Residual components below a minimum area threshold were discarded.

**Wave centerline.** For each wave mask, the centerline was defined as the column-wise mean x-coordinate across all mask-positive rows, smoothed by a moving-average filter (window 200 pixels).

**Wave speed.** A single global speed was computed per experiment as the horizontal displacement between manually annotated start and end coordinates divided by the annotated time span (mm $h^{-1}$). These positions were annotated manually in the experiments, to avoid artifacts caused by irregularities in the automatically detected wave shape, and detected automatically in the simulations.

**Wave bulkiness.** The maximal inscribed radius of each frame's wave mask was computed as the maximum value of the Euclidean distance transform of the mask interior, converted to mm (30 px/mm for experimental data; 20 px/mm for simulation sets), and averaged over the sampled frames of a recording. Note that *wave bulkiness* is our primary lateral measure of wave morphology. Because the front advances non-uniformly, the wave boundary is locally irregular, making cross-sectional width sensitive to the specific sampling location and orientation along the centerline. *Wave bulkiness* (yellow circles in Fig. 4b-i) captures wave extent in a way that is invariant to local shape irregularities, and is therefore a better measure than the standard cross-sectional width (the mean length of sampled normal cross-sections; green lines in Fig. 4b-i) in our case.

##### 1.1 Front kinematics

Front position and width were measured in a separate analysis of the raw image series, independent of the wave mask and of the annotated start and end positions from which *wave speed* is computed (Supplementary Fig. 1). Every tenth frame (100 s) of the wave phase was sampled, skipping its first 100 min or first fifth of its duration, whichever is shorter, while the front forms. Each sampled frame was flat-field corrected, inverted and averaged down each image column within the interior of the food patch, away from its edges, giving a density profile along the direction of advance. Stacked over time, the profiles form the kymographs in Fig. 4b-ii and Supplementary Fig. 1a.

Each frame's density profile was baseline-corrected by subtracting its 10th percentile, then smoothed with a moving average of approximately 1 mm. The leading and trailing edges are the half-maximum crossings ahead of and behind the profile peak, and the front width is the distance between them. Both edges count as detected when they lie inside the measured region and the leading edge passes the 2 mm guard described in the Methods. Front speed is the slope of an ordinary least squares fit of the leading edge positions against time, with $R^2$ from the same fit. Stationarity of the profile is described by the within-replicate coefficient of variation of the front width, the Spearman correlation of width against time, and the fitted percentage change in width across the traverse; the traverse ratio is the distance advanced divided by the mean front width. Values of these quantities for each strain in the experiments and in the simulations are given in Supplementary Data 1.

#### 2. Coverage Geometry Extraction

**Flat-field correction.** Each raw video frame was corrected for non-uniform illumination using a pre-computed flat-field estimate. Corrected frames were converted to double-precision floating point and normalized to [0,1].

**Temporal median filtering and TCI generation.** Corrected frames were sampled at 10-frame intervals (100 s at 0.1 fps). A sliding temporal median was computed over six consecutive sampled frames (~10-minute effective window). The TCI was generated by taking the pixel-wise minimum of these median-filtered values across the entire recording:

$$\mathrm{TCI}(x,y) = \min_t \left[ \mathrm{median}_{s \in \mathrm{window}(t)} \, I(x,y,s) \right]$$

where $I(x,y,s)$ is the corrected intensity at pixel $(x,y)$ at sampled frame $s$. This minimum-of-medians operation retains the darkest sustained signal at each location (corresponding to peak worm density), while the temporal median suppresses transient single-worm events. Parameter choice is validated empirically. See Supplementary Methods, Coverage Geometry Extraction, TCI parameter validation.

**Food patch masking.** For experimental data, the food patch boundary was defined from manually annotated polygon coordinates. For simulation sets, the boundary was extracted directly from simulation geometry. All downstream analyses were restricted to pixels inside this mask.

**Hole segmentation.** Otsu's method was applied to TCI pixel values inside the food-patch mask to produce a binary image. Connected components in the inverted binary image (bright = worm-free regions) were identified. Components were excluded if: (i) their area was below a minimum threshold (100 pixels at 30 px/mm for experimental data, 10 pixels at 20 px/mm for simulation sets), or (ii) more than 15% of their perimeter coincided with the food-patch boundary. Both cutoffs, 0.111 mm² for experiments and 0.025 mm² for simulations, lie well below the *large hole areas* measured (Supplementary Table 2).

**Large hole area (5th rank).** Retained holes were sorted by area in descending order; the 5th-largest area was taken as the metric. Rank stability was validated across $k$ = 1–15 (Supplementary Methods, Coverage Geometry Extraction, Hole-rank metric stability).

**TCI bulkiness (maximal inscribed radius).** The maximum value of the Euclidean distance transform of the dark (worm-visited) foreground mask in the TCI, converted to mm.

##### 2.1 TCI parameter validation

The TCI provides a cumulative spatial record of worm visitation by retaining, for each pixel, the minimum intensity observed across the entire experiment after temporal-median pre-filtering. Below we explain why the minimum is used rather than another summary statistic, and why a temporal median buffer is applied before taking it.

**Minimum versus alternative projections.** Time-averaged projections conflate transiently and persistently visited pixels: a location densely occupied for a short time and one sparsely occupied throughout the experiment can produce the same mean intensity, merging bundles (visited areas) and holes (unvisited areas) into a uniform gray field. Maximum intensity projections highlight the brightest (worm-free) moments, inverting the biologically relevant signal. Minimum projection captures the darkest moment each pixel experienced, corresponding to its peak local worm density. Because worms are semi-transparent in our bright-field microscopy, this minimum directly encodes the most concentrated local visitation, preserving bundled structure and enabling the distinction between persistently visited bundles and under-visited holes that underlies all downstream coverage geometry metrics.

**Temporal median buffering.** Taking a pixel-wise minimum over all raw frames would cause isolated worms or brief imaging artifacts to register as persistent occupancy, artificially filling holes. A sliding temporal median filter, computed over 6 frames sampled every 10 raw frames at 0.1 fps (approximately one frame per 100 s) and spanning a 10-minute window, suppresses these transient single-worm signals while retaining the sustained occupancy signature of the traveling wave. The minimum is then taken over the resulting median-filtered sequence, so only events persisting across the buffer window contribute to the TCI. The choice of parameters is justified empirically in Supplementary Fig. 8.

**Subsamples per 10-minute buffer.** At a single subsample per buffer the TCI is built from one frame every ten minutes and therefore misses some of the regions that worms visited, making holes appear spuriously large and bundles spuriously thin in all three cases. The bias is largest for the N2 (high *TCI bulkiness*) case, where *large hole area* is 3.8-fold overestimated (5.29 vs 1.38 mm² at default) and *TCI bulkiness* is 2.4-fold underestimated (4.06 vs 9.80 mm). From 6 subsamples onward, both metrics plateau for all three cases (maximum change between 6 and 60 subsamples: 18% for *large hole area*, 10% for *TCI bulkiness*). Increasing to 12 or 60 subsamples produces no meaningful additional change, confirming that 6 is not a boundary choice (Supplementary Fig. 8a–b).

**Buffer duration.** At 1-minute buffers the window is too narrow to fully suppress transient worm visits: for the N2 (high *TCI bulkiness*) case, *large hole area* is 2.5-fold underestimated (0.56 vs 1.38 mm² at default), because brief individual worm appearances darken genuine hole pixels in the temporal median, and these artifacts survive the minimum projection. At 5 minutes, values approach but have not yet fully converged to the default. Above 10 minutes, over-aggregation sets in: as the buffer spans frames both before and after the wave has passed a location, the resulting median is brighter than the true wave-passage minimum, effectively erasing the contrast between bundles and holes. Between 10 and 30 minutes, *large hole area* for the same case inflates 9.1-fold (1.38 vs 12.56 mm²) as brightened bundle pixels cross the Otsu threshold and are reclassified as holes, while *TCI bulkiness* collapses 3.7-fold (9.80 vs 2.63 mm) as the worm-visited region contracts correspondingly. The 10-minute default therefore sits between two failure modes: insufficient noise suppression at short durations and temporal over-aggregation at long durations (Supplementary Fig. 8c–d).

Default parameters (6 subsamples, 10-minute buffer) are shown in Supplementary Fig. 8e–g, with the 5th-largest hole and the maximal inscribed radius marked in yellow for each case. The single-subsample failure mode (Supplementary Fig. 8h–j) and the 30-minute-buffer failure mode (Supplementary Fig. 8k–m) illustrate both extremes.

#### 2.2 Hole-rank metric stability

Color-coded rank matrices for all experiments (N2 and *npr-1* combined, $n = 22$), ordered by the 5th-largest hole rank, show a consistent gradient pattern across columns $k = 3$ to $k = 15$, confirming that the relative ordering of experiments is preserved across this range (Supplementary Fig. 9a). In contrast, the $k = 1$ column departs visibly from the gradient, indicating that the largest hole is an unreliable descriptor of a replicate, consistent with its susceptibility to transient imaging artifacts, and motivating the use of $k = 5$ as the representative *large hole area* metric.

Spearman rank correlation between the ordering produced by the 5th-largest hole and that produced by the $k$th-largest hole reaches 0.87 at $k = 3$, 1.0 at $k = 5$ by definition, and remains above 0.94 through $k = 10$, with a gradual decline to 0.91 at $k = 15$. The markedly lower value at $k = 1$ ($\rho = 0.66$) corroborates the rank-matrix pattern (Supplementary Fig. 9b; Supplementary Table 4). Together, these analyses confirm that $k = 5$ sits within a stable regime ($k = 3$–15) and is not a cherry-picked value.

#### 3. Agent-Based Simulation

The model implements the four components named in the main text as follows: (i) body elongation, the ring-buffer segment chain described under Agent body model; (ii) roaming-dwelling switching, the density-mismatch transition rules under Roaming-dwelling switch; (iii) *PD* navigation, the eight-neighbor density gradient rule under Density gradient navigation (movement priority 2); (iv) food chemotaxis, the odor-guided movement rule under Food chemotaxis (movement priority 1). The interaction logic of these components across the spatial arena is described in Section 4. Their dependency structure is shown in Supplementary Fig. 3.

**Spatial domain.** The simulation arena is a 30 x 30 mm square on a 600 x 600 pixel grid (20 px/mm). Agent positions are stored as continuous floating-point coordinates; density calculations use floored integer pixel coordinates. The food patch, a 21 x 12 mm rectangle with rounded corners on the left side, occupies the vertically centered right portion of the arena to mimic the experimental geometry. The worm reservoir occupies an elliptical region on the left side, non-overlapping with the food patch.

**Agent body model.** Each agent is represented as a chain of 20 (x,y) coordinate pairs spaced ~50 µm apart, giving a total body length of ~1 mm and an aspect ratio of 1:20. At each step, a new head coordinate is added in the direction of movement and the rearmost (tail) segment is removed, so the chain advances while keeping body length constant. All 20 segments contribute to local density calculations.

**Round agent variant.** For the leave-one-out simulations without body elongation, agents arrange their 20 segments in a compact circular cluster centered on the head position, preserving total segment number while removing body anisotropy. A short history tail (~3 segments) is preserved to maintain a directional memory vector.

**Roaming-dwelling switch.** Each agent switches stochastically between dwelling and roaming. Per-step transition probabilities depend on a normalized density-mismatch score *rel*, computed as the sum of *distFromOptimal* values across all body-segment pixels, normalized by 20 × *PD*, where *distFromOptimal* = |*local_density* − *PD*| for under-dense pixels and 3 x (*local_density* − *PD*) for over-dense pixels. The transition probabilities are:

For $0 \le rel \le 1$:

$$p_{\text{leave\_roam}} = (1 - rel) \times 1 + rel \times (1/avgRoamingSteps)$$

$$p_{\text{leave\_dwell}} = (1 - rel) \times 0 + rel \times (1/avgDwellingSteps)$$

For $rel > 1$:

$$p_{\text{leave\_roam}} = (1/avgRoamingSteps) \times \exp(-(rel - 1))$$

$$p_{\text{leave\_dwell}} = (1/avgDwellingSteps) + (1 - 1/avgDwellingSteps) \times (1 - \exp(-(rel - 1)))$$

A minimum floor of 0.25x the baseline rate is applied to both probabilities. Initial states are drawn from the stationary distribution of the baseline rates, with one in three agents roaming.

**Food chemotaxis (movement priority 1).** An odor field is computed at each step from the current food patch state using a multi-threshold exponential decay approximation. A roaming agent enters odor-guided mode with probability:

$$p_{\text{guided}} = \sigma((\textit{foodThreshold} - f) / \textit{softness}) \times \sigma((s - \textit{odorThreshold}) / \textit{softness})$$

where σ is the logistic function, *f* is local food concentration, *s* is local odor magnitude, *foodThreshold* = 0.2, *odorThreshold* = 0.4, *softness* = 0.05. An odor-guided agent moves one step in the local gradient direction plus angular noise drawn from uniform (−45°, +45°), clamped to arena bounds.

**Density gradient navigation (movement priority 2).** Roaming agents not in odor-guided mode examine the eight neighboring pixels. The neighbor with density closest to *PD* is the target, with an excess over *PD* penalized three times as much as an equal shortfall. If a density gradient exists (neighbors are not all equal after removing self-overlapping segments), the agent steps toward that target; otherwise the agent continues in the current head-to-tail direction (priority 3: correlated random walk). Angular noise (±45°) is added in all cases.

**Food consumption.** At each pixel with *n* agent heads present:

$$n_{\text{eff}} = 1 + (n - 1) / (1 + (1/\textit{collectiveFeedingEfficiency}) \times (n - 1))$$

Food concentration updates as $F \leftarrow F \times r^{n\text{eff}}$ with $r = 0.999$ and *collectiveFeedingEfficiency* = 0.2.

**Simulation termination.** Simulations were terminated when the residual food quantity fell below 50% of its initial value within the vertical stripe located 5% to 10% from the right edge of the arena. This criterion was selected to optimize the simulation duration relative to patch consumption while minimizing confounding edge effects.

**Video output.** Simulation state is rendered at 0.1 fps. Pixel intensity reflects local worm density, scaled by opacity = 0.1, inverted, and clipped to [0,1] to mimic bright-field microscopy.

**Nematic alignment analysis.** Alignment was measured in separate simulations of 70 agents in a 50 x 50 mm arena at *PD* = 12, 30 min each (120 runs with elongated agents and 1,200 with round agents). Pairwise interaction events were identified as two single-agent trajectories that merged and then separated. Each agent's direction of travel was taken as its displacement over the two body lengths before the meeting point and over the two body lengths after it; the angle between the two agents' directions was recorded before and after the interaction, and a pair that moved on together for two body lengths after meeting was recorded as parallel. For elongated agents, distributions were corrected for geometric sampling bias by the cross-sectional width of two rectangular bodies at each approach angle. For round agents no geometric correction was applied, as circular bodies present an isotropic cross-section. In both cases a non-zero baseline *r* is expected from kinematic encounter bias: in a correlated random walk, head-on encounters occur at relative speed ≈ 2v and are over-represented, while same-direction encounters require one agent to be dwelling while the other overtakes (relative speed ≈ v) and are under-represented; both approach directions collapse to the same nematic angle, producing non-zero *r* independent of any alignment effect. The relevant quantity for assessing nematic alignment induced by interaction is therefore $\Delta r = r_{\text{after}} - r_{\text{before}}$ rather than absolute *r*. Mean resultant length *r*, a standard measure of directional concentration ranging from 0 (uniform) to 1 (perfect alignment), was computed from the corrected distribution of doubled angles, so that parallel and antiparallel pairs count alike.

#### 4. Numerical illustration of component interactions

The three behavioral components (roaming-dwelling switching, *PD* navigation, and food chemotaxis) do not act independently; their interaction produces distinct effective behaviors across the arena that shift dynamically as food is consumed. Three characteristic situations illustrate this.

**Agent inside a cluster at *PD*, away from food.** When local density matches *PD* (normalized mismatch *rel* = 0), the per-step probability of leaving the dwelling state collapses to its minimum floor of 0.0025 per step, a mean dwelling bout of 100 s, while the probability of leaving roaming reaches 1. A roaming agent whose segments sit at *PD* therefore dwells at the next step (0.25 s). Among the small roaming fraction permitted by the floor, chemotaxis activates with a probability that rises with odor magnitude: the odor field decays exponentially with distance d from the food boundary ($\beta \approx 0.17$ mm$^{-1}$), reaching the 50% activation threshold at $d^* \approx 5.4$ mm and falling below 10% beyond ~7 mm. At the reservoir center (~5–6 mm from food), an agent in a cluster therefore experiences: $p$(dwell this step) ≈ 0.9975, $p$(chemotaxis | roaming) ≈ 0.5, yielding an effective per-step probability of moving toward food of roughly 0.0025 x 0.5 ≈ 0.13%. The cluster is essentially static.

**Agent in the front on intact food.** A cluster arriving at a food-rich location remains near *PD* (dwells for the same reason as above). Chemotaxis is additionally suppressed by the food availability factor of $p_{guided}$, implemented as a logistic function that approaches zero when local food concentration exceeds ~0.2 (normalized): at fully intact food levels (~1.0), the odor ≈ 0.018, reducing $p_{guided}$ by ~50-fold regardless of odor strength. Agents on intact food therefore have an effective chemotaxis probability of 0.0025 x 0.018 x the odor factor, at most 0.0045%, effectively zero. They dwell and eat.

**Agent in the front as food depletes.** Food consumption reduces local food concentration progressively at each step. As the local value falls below 0.2, the food availability factor rises above 0.5 and approaches 1 at full depletion. Simultaneously, the odor field, recomputed each step from the current food distribution, now carries a gradient pointing toward intact food ahead. The floor-rate roaming trickle remains the only eligible pool (0.25% of steps) but among those roaming steps, chemotaxis guidance now rises steeply: at 50% local depletion and 5 mm from the food edge, $p_{guided}$ ≈ 0.0025 x 0.5 x 0.5 ≈ 0.06% per step, and at full depletion $p_{guided}$ ≈ 0.0025 x 0.98 x 0.5 ≈ 0.12%. This gentle but consistent bias draws the small roaming fraction toward the receding food edge, pulling the cluster forward gradually as food disappears beneath it. The wave therefore advances not by agents breaking away from the cluster, but by the cluster being gently steered forward at floor-rate by the combined release of chemotaxis suppression and the shifting odor gradient.

The behavioral regimes across the arena with cluster stasis, food-pinned dwelling, and depletion-gated forward pull, are therefore not pre-specified spatial zones but emerge dynamically from the interaction of the three components with the evolving food distribution.

**Effective speed and bout durations.** Agents dwell most at *PD* and less the further local density departs from it, so on average they move slowest at *PD*. This effective speed follows from the switching rule alone: it is the roaming speed $v_{roam}$ (0.2 mm $s^{-1}$; Supplementary Table 3) multiplied by the fraction of time spent roaming,

$$v(\rho) = v_{roam} \times p_{leave\_dwell} / (p_{leave\_roam} + p_{leave\_dwell})$$

where $p_{leave\_dwell}$ and $p_{leave\_roam}$ are the per-step probabilities of leaving dwelling and roaming at local density ρ (Section 3); the mean bout in each state is the reciprocal of its leave probability. Effective speed is 1.8 mm $h^{-1}$ at *PD*, against about 20 mm $h^{-1}$ at a quarter of *PD* and 716 mm $h^{-1}$ at twice *PD*, and is the same function of ρ/*PD* for both preferred densities. Below *PD* it falls steeply enough with increasing density to meet the classical criterion for a motility-induced instability, $d \ln v / d \ln \rho < -1$, between 0.15 *PD* and 0.75 *PD* (−1.9 at 0.5 *PD*); for *PD* = 6 the band starts at ρ = 1, the lowest density an agent can read, which is 0.17 *PD*. Above *PD*, *v* rises with density.

The baseline bouts of Supplementary Table 3 (25 s dwelling, 12.5 s roaming) are realized only at 4/3 *PD*. Dwelling and roaming bouts are equal, 12.6 s each, at about 1.34 *PD*; below that density dwelling bouts are longer than roaming bouts. Mean bouts are 33 s dwelling and 0.9 s roaming at a quarter of *PD*, 100 s and 0.25 s respectively at *PD*, and 0.3 s and 50 s respectively at twice *PD*.

#### 5. Leave-one-out simulations

Figure 3 shows the four leave-one-out simulations, in which a single model component is removed while all other parameters remain as listed in Supplementary Table 3 (1,500 agents, default rectangular arena and food geometry). Supplementary Movie 4 shows the full dynamics of all four conditions side by side.

**Body elongation (Fig. 3a,b; Supplementary Movie 4a).** Round-body agents retain the same 20 body segments and all identical parameters as elongated agents. The sole change is how the segments are spatially arranged: instead of a chain forming a 1:20 aspect-ratio body, segments are distributed in a compact circular cluster around the head. This removes the geometric anisotropy responsible for nematic alignment without altering agent number, density sensing, roaming-dwelling switching, or food chemotaxis. Angular distribution analysis of separate alignment runs (Section 3, Nematic alignment analysis) confirmed that pairwise interactions increased alignment exclusively for elongated agents. For these agents, the mean resultant length increased from $r = 0.11$ before interactions to $r = 0.45$ after ($\Delta r = 0.34$). In contrast, round agents showed no increase ($r = 0.20$ before, 0.15 after; $\Delta r = -0.05$), consistent with an absence of interaction-induced alignment. The non-zero baseline ($r = 0.20$) for round agents reflects a kinematic encounter bias from correlated locomotion rather than true nematic alignment, as head-on encounters were over-represented relative to same-direction meetings. Without elongation, agents consume the food patch from all directions in a uniform advancing front with no cryptic network.

**Roaming-dwelling switching (Fig. 3c; Supplementary Movie 4b).** The dwelling state is disabled so agents transition immediately back to roaming upon stopping. Roaming-only agents initially produce a front and a transient cryptic network, but these collective structures subsequently fragment into randomly distributed short-lived aggregates. The dwelling state, and the local density amplification it provides, is therefore necessary for sustaining the cryptic network.

***PD* navigation (Fig. 3d; Supplementary Movie 4c).** *PD* navigation is disabled so all locations are treated as equally attractive regardless of local worm density. Agents navigate by chemotaxis alone. The resulting TCI shows agents distributed approximately uniformly across the food patch, with a slight intensity increase near the patch boundaries, and no cryptic network, confirming that density-dependent aggregation is necessary for bundle and hole formation.

**Food chemotaxis (Fig. 3e; Supplementary Movie 4d).** The food chemotaxis component is disabled: the food gradient is computed but agents do not navigate toward it. Agents form continuously remodeling visible networks but no traveling wave, confirming that directed chemotaxis, not merely food presence, is required for wave formation. Notably, when chemotaxis is removed but *PD* is varied between strains (Supplementary Movie 4d shows N2 *PD* = 6 left, *npr-1 PD* = 12 right), the visible networks still differ: the higher *PD* of the simulated *npr-1* produces denser, more interconnected aggregates despite identical food environments.

#### 6. Sensitivity analysis

The sensitivity analysis varied body length (the number of body segments), the baseline roaming and dwelling bouts, odor threshold, angular noise and collective feeding efficiency (Supplementary Table 3); *PD* was not perturbed, but its values in the two simulated strains differ twofold and both produce the wave and its cryptic network (Fig. 4). Each parameter was set to 0.8x and 1.2x its default with all others held at default, giving twelve perturbed conditions and one shared default condition. Every condition was run as 40 independent replicates at 1,500 agents and *PD* = 12, and all 520 runs were processed through the same analysis pipeline as the simulated strain sets (Methods, Simulation Analysis). Survival of the wave and of the cryptic network was judged from the metrics, which were obtained in all 40 replicates of every condition and whose condition means stayed within 39% of the default (Supplementary Table 1), and from TCIs of the conditions that displace coverage geometry the furthest (Supplementary Fig. 5b).

A response was flagged when $|z| > 2$, where $z$ is Welch's $t$ statistic, the difference between the perturbed and default means divided by its standard error (Supplementary Table 1). The principal axis in Figure 5b is the first principal component of the twelve conditions' percentage deviations from the default in *large hole area* and *TCI bulkiness*, computed about the origin, the default condition, rather than about their mean.

#### 7. Independence of the metrics from wave phase duration

Wave phase durations ranged from 90 to 335 min. To rule out duration as a confound, we computed the Spearman correlation between duration and each metric separately per strain (Supplementary Fig. 7; Supplementary Table 4; N2 $n$ = 14, *npr-1* $n$ = 8). Coverage geometry metrics (*large hole area*, *TCI bulkiness*) showed no significant correlation with duration in either strain. One case is borderline, *large hole area* in N2 ($\rho$ = 0.52, $P$ = 0.059), with no counterpart in *npr-1* ($\rho$ = 0.14, $P$ = 0.752). *Wave bulkiness* showed no significant correlation with duration in either strain. *Wave speed* is shown for completeness only: a slower wave takes longer to cross the patch, so a negative association with duration is expected, and both strains show a negative trend (Supplementary Fig. 7a). Because wave phase durations were statistically indistinguishable between strains (Methods, Wave phase duration), duration does not confound the between-strain comparisons reported in the main text.

### Supplementary Notes

#### Supplementary Note 1. Biological realism of model parameters and acknowledgement of quantitative limitations

The simulation was designed to identify the components sufficient to generate the observed collective dynamics and relative between-strain differences, rather than to reproduce absolute metric values.

**Parameter grounding.** Body dimensions (1 mm total length across 20 segments, ~50 μm width per segment) match published adult *C. elegans* measurements[1]. The integration time step ($\Delta t$ = 0.25 s) is set so that an agent moving at typical roaming speed (four segments, or 0.2 mm, per second) displaces one segment per step. While simplified for simulation step-size consistency, our assigned baseline durations for dwelling and roaming (25 s and 12.5 s) qualitatively align with the behavioral partitioning described in published ethograms on bacterial lawns, where the dwelling state is the predominant phase[3,4]. These durations are the reciprocals of the baseline switching rates; realized bouts vary with local density (Supplementary Methods, Numerical illustration of component interactions, Effective speed and bout durations). The strain-specific value of *PD* (6 segments/pixel for N2; 12 for *npr-1*) reflects the enhanced aggregation phenotype conferred by the *npr-1(ad609)* null mutation; its value was determined by matching observed wave morphology rather than by independent measurement. All parameters are listed in Supplementary Table 3.

**Scale of quantitative discrepancy.** Absolute metric values in the simulations differ from experiments by up to approximately one order of magnitude across metrics (Supplementary Table 2). For N2 and *npr-1* respectively, the simulated values are lower by factors of 6.5 and 5.3 for *wave speed*, 6.1 and 5.5 for *wave bulkiness*, 7.3 and 1.6 for *large hole area* (2.7 and 1.3 in hole diameter), and 9.6 and 5.8 for *TCI bulkiness*.

**Interpretation.** We prioritized identifying the components sufficient to generate the observed behavioral repertoire rather than fitting model parameters to collective-level outputs. Because none of the metrics (*wave speed*, *wave bulkiness*, *TCI bulkiness*, *large hole area*, or the direction of the strain differences) was used to constrain the parameters, the agreement between model and experiment reflects the explanatory reach of the four identified model components rather than parameter absorption. Systematic calibration to absolute values would be a valuable next step: it could identify which specific aspects (e.g. unreported density-sensing thresholds, biophysical constraints on body mechanics, or food-patch geometry differences) account for the quantitative gap, and in doing so would deepen mechanistic understanding beyond what component-level identification alone achieves. Whether the quantitative gap between experiments and simulations reflects a missing mechanism or an unexplored parameter region remains an open question. The simulation reproduces the visible network, the cryptic network and full wave phenomenology, and the direction and statistical significance of strain differences in both wave metrics. The model also contains a trade-off between the two coverage geometry metrics, in line with the spread of N2 experimental replicates, but not what causes that spread (Fig. 5; Supplementary Note 2). Taken together, these features support that within the model, the four components are sufficient to generate both collective phases of the wurmuration behavior.

#### Supplementary Note 2. Open questions and directions for future investigation

Three questions remain open for future investigations.

**Network-to-wave transition mechanism.** The transition between wurmuration phases (reservoir drainage, wave initiation, and eventual collective dissolution) is characterized here at the macroscale through TCIs, wave metrics and front kinematics. The individual-level dynamics of this transition remain uncharacterized: whether the drainage of the visible network into the wave is triggered by a threshold in local food odor, a critical reduction in reservoir density, or a temporal accumulation of individual chemotaxis-driven departures is not resolved by the current experimental data. Multi-worm tracking at population scale across both phases could test the latter two of these explanations and show whether the transition is a clear switch or a gradual process.

**Sources of variability between N2 experimental replicates.** In the sensitivity analysis, parameter perturbations move the two coverage geometry metrics along a trade-off (Fig. 5b; Supplementary Table 1). This trade-off in the model is in line with the trend across N2 experimental replicates, in which the two metrics are negatively though not significantly associated (Fig. 5a). Variation between replicates in the fraction of worms joining the collective movement could be one variable that shifts N2 replicates along the model's trade-off, an explanation that remains to be tested in the future.

**Model calibration.** The simulations reproduce directional trends and relative strain differences in both wave metrics but not in coverage geometry, and differ from absolute experimental values by up to approximately one order of magnitude (Supplementary Note 1). Closing this gap requires independent measurement of the biological quantities the model treats as free parameters: the density deviation thresholds governing roaming-dwelling transition probabilities, the spatial range of food odor sensing, and the effective density footprint of a single worm's body. *PD* itself was fitted rather than measured (Supplementary Note 1). Whether experimentally raising density preference in the worms makes the wave slower and more compact, as raising *PD* does in the model (Fig. 4a), remains to be tested.

### Supplementary References


1. Brenner, S. The genetics of Caenorhabditis elegans. Genetics 77, 71–94 (1974).
2. Ding, S. S., Schumacher, L. J., Javer, A. E., Endres, R. G. & Brown, A. E. Shared behavioral mechanisms underlie C. elegans aggregation and swarming. eLife 8, e43318 (2019).
3. Arous, J. B., Laffont, S. & Chatenay, D. Molecular and Sensory Basis of a Food Related Two-State Behavior in C. elegans. PLOS ONE 4, e7584 (2009).
4. Flavell, S. W. et al. Serotonin and the Neuropeptide PDF Initiate and Extend Opposing Behavioral States in C. elegans. Cell 154, 1023–1035 (2013).
